# BET protein inhibitor JQ1 reduces inflammation and hippocampal amyloid-β level without altering Tau phosphorylation in LPS-challenged adult wild-type mice

**DOI:** 10.1101/2025.09.22.677813

**Authors:** Marta Matuszewska, Magdalena Cieślik, Dorota Sulejczak, Anna Wilkaniec, Grzegorz A. Czapski

## Abstract

**Introduction:** A growing body of evidence highlights the role of infection and inflammation in the progression of Alzheimer’s disease (AD). In this study, we aimed to analyze the impact of JQ1, an inhibitor of bromodomain and extraterminal domain (BET) proteins, which are key readers of the epigenetic acetylation code, on AD-related gene expression changes and biochemical alterations in the hippocampus during a lipopolysaccharide (LPS)-induced systemic inflammatory response in mice.

**Methods:** JQ1 and LPS were administered intraperitoneally to adult male wild-type C57BL/6J mice. Changes in selected general and brain-specific parameters were measured for up to 12 h.

**Results:** Our results demonstrated that inhibition of BET proteins reduced LPS-induced sickness behavior and time-dependent elevation of proinflammatory signaling. LPS did not significantly alter amyloid-β (Aβ) levels; however, a significant reduction in Aβ load was observed in JQ1-treated animals overall, suggesting that BET proteins play a crucial role in regulating Aβ levels in the brain. At the same time, JQ1 treatment did not affect LPS-induced increases in phospho-Tau levels.

**Discussion:** Our results suggest that inhibiting BET proteins, in addition to their anti-inflammatory action, may be an effective strategy for reducing Aβ levels in the brain. However, a mechanistic explanation of this phenomenon requires further investigation.

## Introduction

Alzheimer’s disease (AD) is the most prevalent neurodegenerative disorder and a leading cause of dementia in the aging population. The classical neuropathological hallmarks of AD are extracellular amyloid-β (Aβ) aggregates (senile plaques) and intracellular neurofibrillary tangles (NFTs), which are composed of hyperphosphorylated Tau protein. Early-onset AD (EOAD), accounting for less than 5% of all cases, is caused by mutations in the genes encoding presenilin 1 (PSEN1), presenilin 2 (PSEN2), and amyloid-β (Aβ) precursor protein (APP), resulting in accelerated Aβ deposition (Chartier-Harlin et al., 1991; Sherrington et al., 1995). In contrast, the vast majority of cases are sporadic, late-onset AD (LOAD), wherein complex interactions among genetic susceptibility, environmental factors, and immune responses contribute to disease onset and progression.

The amyloid cascade hypothesis, which postulates Aβ accumulation as the primary driver of neurodegeneration, has been increasingly challenged by numerous unsuccessful clinical trials targeting Aβ (Armstrong, 2014; Kepp et al., 2023; Weinstock, 2024). Even though anti-Aβ antibody-based therapy was finally approved (Jucker and Walker, 2023), its modest benefits and numerous disadvantages highlight the need to explore alternative hypotheses. Growing evidence indicates that neuroinflammation is not merely a bystander but rather a prerequisite for neurodegeneration in AD. For example, cognitively resilient individuals with high Aβ and Tau loads exhibit reduced neuroinflammatory markers, highlighting the modulatory role of inflammation in AD susceptibility (Barroeta-Espar et al., 2019; Perez-Nievas et al., 2013).

Systemic inflammation, such as that induced by bacterial or viral infections, is a well-established risk factor for accelerated AD progression (Bu et al., 2015; Douros et al., 2021; Sun et al., 2022b). Early observations suggested a detrimental role for the inflammation-induced release of potentially neurotoxic mediators (McGeer and McGeer, 1998). Peripheral administration of lipopolysaccharide (LPS), a potent endotoxin derived from Gram-negative bacteria, triggers robust microglial activation, cytokine release, blood–brain barrier (BBB) dysfunction, and transcriptional alterations in the brain (Czapski et al., 2016; Ifuku et al., 2012; Wang et al., 2018b). Rodent studies have demonstrated that both acute and chronic LPS exposure lead to AD-like neuropathology, including Aβ accumulation and Tau hyperphosphorylation (Czapski et al., 2016; Hossain et al., 2018; Ifuku et al., 2012; Kahn et al., 2012; Lee et al., 2008; Wang et al., 2018b), and that these effects may persist long after the initial immune challenge (Wang et al., 2020). Indeed, chronic neuroinflammation and progressive neurodegeneration have been observed following systemic LPS administration in mice (Holmes et al., 2009), with analogous mechanisms implicated in human AD progression (Cunningham et al., 2009).

Although AD is characterized by slow progression, acute systemic inflammation may act as a mechanistic “first hit,” initiating or amplifying neurodegenerative cascades (Cunningham et al., 2009). This notion is supported by evidence that systemic LPS challenges induce lasting cognitive impairments and neuroinflammatory reprogramming (Qin et al., 2007; Teeling and Perry, 2009). While such acute models in young, wild-type mice do not fully recapitulate the complexity of AD, they are valuable for dissecting early immune–brain interactions and identifying potential therapeutic targets (Cunningham et al., 2009; Lee et al., 2008).

One proposed mechanism involves CD33, a microglial immunomodulatory receptor that inhibits the phagocytosis of Aβ. Genetic variants of CD33 are associated with LOAD (Griciuc et al., 2013), and its upregulation has been linked to increased Aβ burden and impaired microglial clearance. Our recent findings demonstrate that systemic LPS elevates hippocampal Cd33 expression in mice (Czapski et al., 2024), suggesting that recurrent peripheral inflammation may potentiate amyloid pathology via this axis.

Epigenetic regulation plays a critical role in orchestrating these neuroimmune responses. Bromodomain and extraterminal domain (BET) proteins—including Brd2, Brd3, and Brd4—function as readers of acetylated histones and interact with transcription factors such as NF-κB to regulate the expression of proinflammatory genes (Martella et al., 2023; Wang et al., 2023). BET dysregulation has been implicated in LPS-induced neuroinflammation (Wang et al., 2020), and BET inhibition using the selective inhibitor JQ1 has demonstrated potent anti-inflammatory effects in both peripheral immune cells and the central nervous system (Belkina et al., 2013; Gilan et al., 2020; Liu et al., 2021; Wang et al., 2020; Wang et al., 2018a; Wienerroither et al., 2014). Notably, JQ1 treatment downregulates Cd33 expression, suppresses LPS-induced cytokine production, and impairs microglial phagocytosis in vitro (Czapski et al., 2024; Matuszewska et al., 2022). Furthermore, BET proteins have been implicated in neuronal transcriptional regulation and memory processes (Korb et al., 2015), suggesting that BET inhibition may confer both immunomodulatory and neuroprotective benefits relevant to early AD. While this acute inflammatory model does not replicate the full chronicity and complexity of human AD, it provides a controlled framework to elucidate initial molecular and epigenetic events that may prime the brain for later neurodegenerative processes.

Given these findings, we aimed to investigate the effects of a single JQ1 administration on neuroinflammatory and AD-related markers in young mice subjected to systemic LPS-induced inflammation. This approach facilitates the study of early transcriptional and epigenetic responses that may underlie the initiation of AD-like pathology and offers insights into the therapeutic potential of BET inhibitors in disease prevention.

## Material and methods

### Animals

Experiments were conducted on 3-month-old male C57BL/6J mice supplied by the Animal House of the Mossakowski Medical Research Institute, Polish Academy of Sciences (Warsaw, Poland). A total of 72 animals were randomly allocated into eight equal groups. No exclusion criteria were established a priori. The animals were maintained under standard conditions, with controlled temperature (22°C ± 10%) and humidity (55% ± 10%). All animal experiments were approved by the II Local Ethics Committee for Animal Experimentation in Warsaw (permission WAW2/060/2020). This study was performed in accordance with the relevant guidelines and regulations. All efforts were made to minimize animal suffering and to reduce the number of animals. All manipulations were performed gently and promptly to reduce stress.

### Experimental design

The JQ1 solution was prepared as described previously (Jostes et al., 2017). Briefly, it was dissolved in DMSO and mixed at a 1:10 ratio with a 10% solution of 2-hydroxypropyl-β-cyclodextrin. LPS (from *E. coli* serotype O55:B5; toxicity 1.5×10^7^ EU/mg; Sigma, St. Louis, MO, USA) was dissolved in saline.

All the treatments were performed in a random order in the morning. Animals (three-month-old males) were transported to the laboratory room and, after 30 min acclimatisation, intraperitoneally injected with JQ1 (50 mg/kg b.w. or 300 µL of the vehicle in respective groups) and, 30 min later, with LPS (i.p.; 1 mg/kg or 50 µL of the vehicle in respective groups) (Belkina et al., 2013). Sickness behavior was estimated every 3 h. After 3 or 12 h, the animals were anesthetized by isoflurane inhalation and decapitated (Fig. 1a).

**Figure 1:**
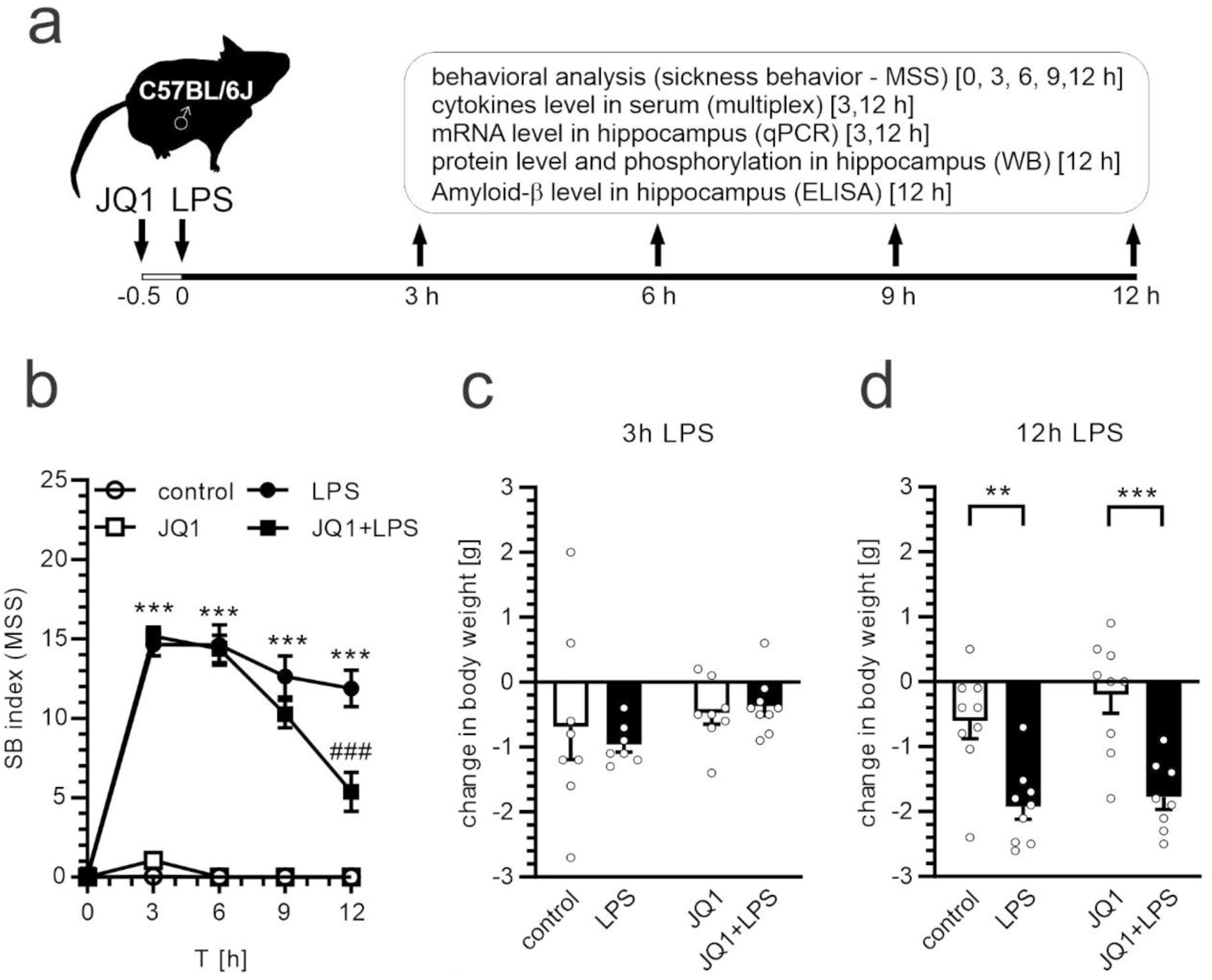
The effect of JQ1 on sickness behavior after LPS-evoked systemic inflammation. (a) Experimental design. Mice were injected with JQ1 (50 mg/kg b.w.) and after 30 min with LPS (1 mg/kg b.w.). Each group received corresponding volumes of vehicles. Sickness behavior was analyzed up to 12 hours post-injection. The level of cytokines in serum and mRNA level in the hippocampus was analyzed 3 and 12 h post-injection. The Aβ load was determined 12 h post-injection. (b) Sickness behavior (SB) index (MSS) was determined every 3 hours after administration of LPS until the end of the experiment. The total body weight change was determined at 3 h (c) and 12 h (d) after administration of LPS. **,*** p<0.01 and 0.001, comparing to respective group: corresponding time point control in (b) or bracket-indicated group in (d). ### p<0.001, comparing to respective (corresponding time point) LPS-treated group, n=8-17 (b) and n=7-9 (c and d). Statistical analyses were conducted using one-way ANOVA followed by the Bonferroni post hoc test.

### Analysis of sickness behavior

The severity of sickness behavior was estimated using a simplified Murine Sepsis Score (MSS) (Shrum et al., 2014). In brief, the animals were observed by an investigator blinded to the treatment groups, and changes in appearance, level of consciousness, activity, response to stimuli, and eye condition were ranked using a scale of 0 to 4 (0 representing normal, 4 indicating severe alteration). In contrast to the original MSS analysis protocol described by Shrum et al. (36), we omitted the assessment of respiration rate and respiration quality due to interpretation challenges.

### Analysis of serum cytokine levels (multiplexing assay)

Blood was collected at 3 or 12 h after LPS administration. Immediately after clot formation, the samples were centrifuged at 1000 × g for 5 min to separate the serum, which was then immediately frozen and stored at -85° C until analysis. Serum cytokine levels were determined using a Bio-Plex Pro Mouse Cytokine 23 assay kit following the manufacturer’s instructions (Bio-Rad) using a Luminex-200 apparatus, before the analysis, the samples were diluted four fold with a dedicated sample diluent.

### Analysis of mRNA levels (qPCR)

Total RNA was extracted from mouse hippocampi using TRI reagent (Thermo Fisher Scientific, Inc.) according to the manufacturer’s protocol. RNA quality and yield were measured using a NanoDrop 2000 spectrophotometer (Thermo Fisher Scientific, Inc.). Potential DNA contamination was digested using DNase I (Sigma-Aldrich) according to the manufacturer’s protocol. Reverse transcription was performed using a High-Capacity cDNA Reverse Transcription Kit with RNase Inhibitor (Thermo Fisher Scientific, Inc.) according to the manufacturer’s protocol. The cDNA was stored at -20°C until use. Quantitative PCR was performed on Applied Biosystems 7500 Real-Time PCR System by using pre-developed TaqMan Gene Expression Assays *Arg1* (Mm00475988_m1), *Abca7* (Mm00442646_m1), *Bin1* (Mm00437457_m1), *Cd2ap* (Mm00815310_s1), *Clu* (Mm01197002_m1), *Cr1l* (Mm00785297_s1), *Il1b* (Mm00434228_m1), *Il6* (Mm00446190_m1), *Nos2* (Mm00440502_m1), *Picalm* (Mm00525455_m1), *Rin3* (Mm00617220_m1), *Tnf* (Mm00443258_m1), *Trem2* (Mm04209424_m1), *Zyx* (Mm00496120_m1). Relative mRNA levels were calculated using the ΔΔCt method, with *Hprt* (Mm00446968_m1) as a reference gene.

### Analysis of protein level and phosphorylation (Western blotting)

Protein immunoreactivity analysis was performed as previously described with modifications (Gąssowska-Dobrowolska et al., 2021). Briefly, tissue samples were homogenized in RIPA buffer, and the protein concentration was determined using a BCA Protein Assay Kit (Thermo Fisher Scientific), with bovine serum albumin as a standard. Samples were mixed with Laemmli buffer and heated at 95°C for 5 min. After SDS-PAGE, the proteins were transferred to a nitrocellulose membrane under standard conditions and used for immunochemical analysis, followed by chemiluminescent detection. Densitometric analysis was performed using TotalLab4 software (NonLinear Dynamics Ltd., Newcastle upon Tyne, UK) using size-marker-based verification. Glyceraldehyde-3-phosphate dehydrogenase (GAPDH) was used to normalize the data. The anti-Tau antibodies were from Santa Cruz Biotechnology (Dallas, TX, USA). The rabbit anti-pTau(Ser199/202) antibodies and rabbit anti-GAPDH antibodies were from Sigma Aldrich (St. Louis, MO, USA). The mouse anti-pTau(Ser396) and rabbit anti-pTau(Ser416) were from Cell Signalling Technology (Danvers, MA, USA). The secondary antibodies used were anti-mouse IgG (GE Healthcare) and anti-rabbit IgG (Sigma-Aldrich). The Tau phosphorylation sites were identified according to the 2N4R isoform (441 kDa) sequence. Details of Western blotting reagents and conditions are presented in Supplementary Table 1.

### Analysis of BET proteins level (ELISA)

Brd2, Brd3, and Brd4 levels were quantitatively determined using an enzyme-linked immunosorbent assay (ELISA) kit (Abbexa Ltd. Cambridge, UK). Briefly, the mouse hippocampi were rinsed with ice-cold PBS to remove excess blood. The tissues were finely chopped and homogenized on ice in PBS using a tissue homogenizer, and the samples were sonicated. The homogenates were centrifuged at 10,000 × g for 5 min and the supernatants were collected. Protein concentration was determined using the BCA assay with bovine serum albumin as the standard. The samples were diluted to a concentration within the acceptable range of 0.01 mol/L in PBS (pH 7.2). Fresh samples were used to prevent protein degradation and denaturation. The ELISA was performed according to the manufacturer’s instructions. The optical density was measured spectrophotometrically at 450 nm using a Multiskan GO Microplate Spectrophotometer (Thermo Fisher Scientific, Inc.). The results were normalized to protein levels.

### Analysis of Amyloid-β level (ELISA)

Quantitative determination of mouse Aβ_1-40_ and Aβ_1-42_ levels was performed using commercially available ELISA kits (Thermo Fisher Scientific, Inc.). Briefly, mouse hippocampi were homogenized in cold 5 M guanidine-HCl/50 mM Tris (pH 8.0). Homogenates were placed in a laboratory rocker at room temperature for 4 h. Then, the samples were diluted tenfold with cold PBS containing a 1× protease inhibitor cocktail and centrifuged at 16,000 × g for 20 min at 4°C. The supernatant was transferred to clean microcentrifuge tubes and used for further analysis, according to the manufacturer’s protocol. The results were normalized to protein levels. The entire assay, including the standard curve, was completed within one day.

### Curated Analysis of Published mRNA Expression Data

In July 2024, the NCBI GEO database (Edgar et al., 2002) was searched for data concerning the impact of JQ1 on human gene expression. Data deposited by Baek and co-workers (Baek et al., 2021), accessible through the GEO Series accession number GSE155408 (https://www.ncbi.nlm.nih.gov/geo/query/acc.cgi?acc=GSE155408), were utilized for a curated analysis of selected genes.

### Statistical Analysis

The group size was calculated with G*Power software, which calculates the minimum required group size based on the size of the Cohen d effect. Standard assumptions adopted: test power = 0.95, significance level = 0.01, groups of equal size. Based on our results from previous experiments with LPS we obtained desired n=9. Statistical analyses were performed using GraphPad Prism version 8.3.0 (GraphPad Software, San Diego, CA, USA). The distribution of data was analyzed using the Shapiro-Wilk test. Outliers were determined using Grubs test and removed if p<0.05. The results are expressed as mean ± SEM. Depending on the experimental design, data were analyzed using Student’s t-test or one-way analysis of variance (ANOVA) with Bonferroni post-hoc test with correction for multiple comparisons. Statistical significance was set at P < 0.05. N refers to independent samples (i.e., various animals).

## Results

Intraperitoneal administration of lipopolysaccharide (LPS) is a commonly used experimental method for inducing a systemic immune response. This response encompasses acute innate mechanisms in the peripheral tissues and central nervous system. In the current study, we administered LPS at 1 mg/kg b.w. (1.5×10^7^ U/kg b.w.) (Czapski et al., 2013; Czapski et al., 2007; Czapski et al., 2006; Czapski et al., 2004; Czapski et al., 2010; Czapski et al., 2016; Jacewicz et al., 2009). This dose evoked a rapid, substantial, and transient response, which was significantly lower than the lethal dose for mice (approximately 50 mg/kg body weight) (Back et al., 2000; Li et al., 2018). However, this dose is high enough to induce a broad range of brain pathologies, including oxidative stress, followed by the activation of PARP-dependent AIF-mediated necroptosis, changes in kinase activity, or inflammatory signaling(Czapski et al., 2013; Czapski et al., 2007; Czapski et al., 2006; Czapski et al., 2010; Czapski et al., 2016; Czapski et al., 2020). To evaluate whether a single administration of a BET protein inhibitor modulates the systemic and central inflammatory responses induced by LPS, we examined a range of peripheral and central variables. At the behavioral level, the response to LPS-evoked endotoxemia is described as a sickness behavior, which includes changes in body temperature, apathy, loss of social interest, and lack of appetite (Harden et al., 2015). We implemented a murine sepsis score (MSS) assessment for semi-quantitative determination of sickness behavior (Shrum et al., 2014). As presented in Fig. 1b, LPS induced a rapid and persistent increase in the level of MSS, which began 3 h after injection and was observed until the end of the experiment, 12 h after exposure (all components of MSS are presented separately in Supplementary Fig. 1). In addition, LPS induced a decrease in body weight, which was indistinct 3 h after injection (Fig. 1c) but was evident at 12 h (Fig. 1d). In animals pretreated with JQ1, the increase in MSS was attenuated and tended to decrease. However, JQ1 did not affect the LPS-evoked changes in body weight.

Although the anti-inflammatory properties of JQ1 are well established, its effects on the levels of specific inflammatory cytokines in blood serum have not been thoroughly investigated. Therefore, we used a multiplexing method to simultaneously analyze the levels of 23 inflammation-related signaling compounds in the mouse serum. Given that no significant changes in the examined parameters were observed in the statistical analysis between the 3-hour and 12-hour time points in the control group receiving the vehicle, we decided to combine these groups into a single unified control group. As expected, LPS induced a rapid increase in the serum levels of nearly all the 23 cytokines tested (Fig. 2). Specifically, the levels of 22 cytokines were elevated at 3 h post-administration and 21 remained elevated at 12 h. Applying the BET protein inhibitor to animals that were not challenged with LPS did not affect their serum cytokine/chemokine profile. However, in the LPS group, JQ1 pretreatment resulted in the augmentation of IL-1α, IL-2, IL-12p70, GM-CSF, and RANTES at 3 h post-LPS, and IL-5, Il-6, IL12p40, Eotaxin, MCP-1, and RANTES at 12 h post-LPS. These results indicate that the BET protein inhibitor had a highly selective effect on the expression of genes related to inflammation. This finding suggests that the mechanism by which JQ1 reduces sickness behavior may involve more than just a reduction in cytokine levels. The complete dataset used to generate the heat map is presented in Supplementary Table 2. The levels of all cytokines exhibited a strong positive correlation with either MSS or with each other (Supplementary Fig. 2).

**Figure 2:**
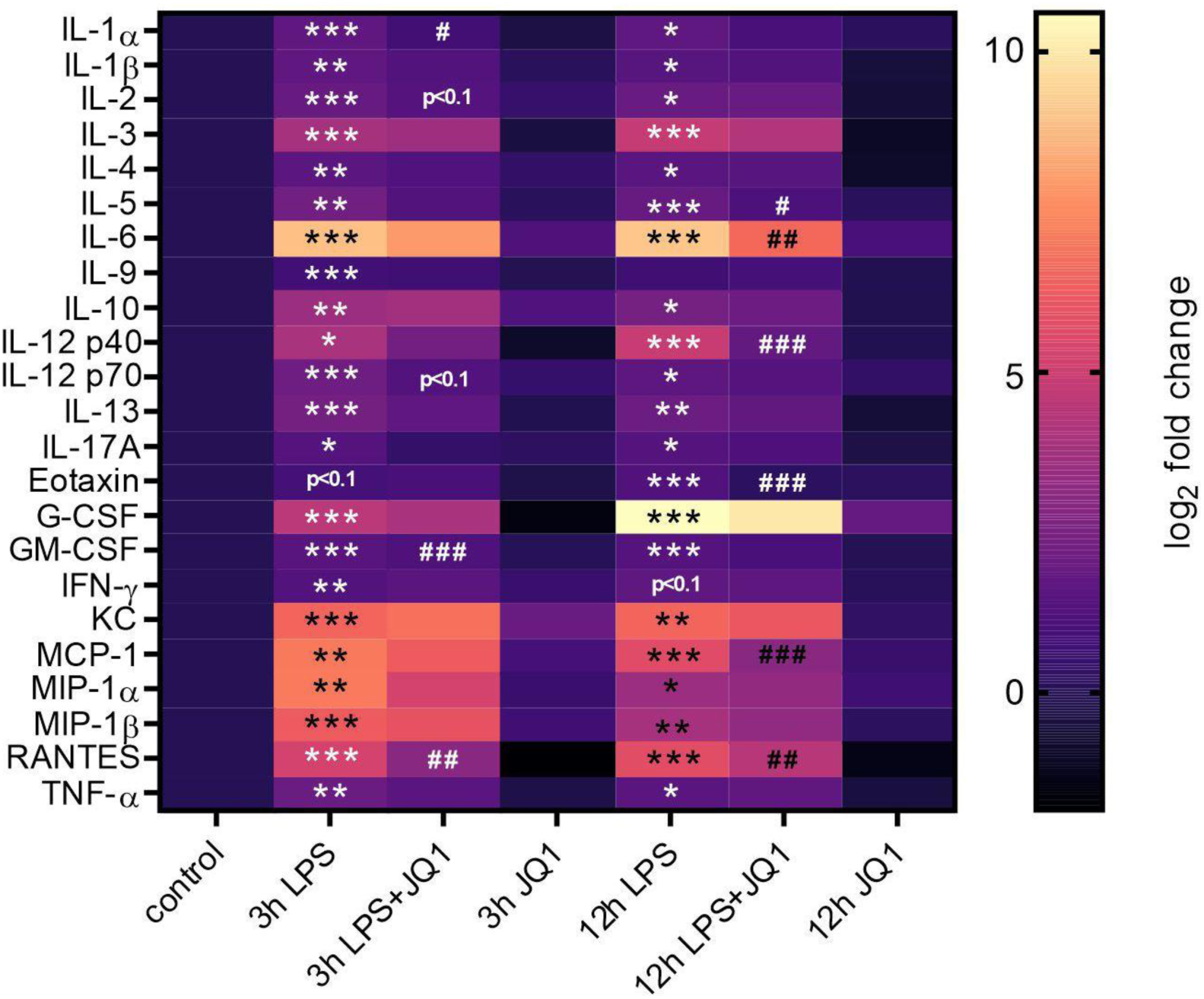
The effect of JQ1 on cytokine levels in blood serum after LPS-evoked systemic inflammation. Mice were injected with JQ1 (50 mg/kg body weight) and after 30 min with LPS (1 mg/kg body weight). The respective groups received the respective vehicle volumes. Cytokine levels in blood serum were determined using the multiplexing method and are presented as a heatmap. The control groups at 3 and 12 h post-injection did not differ statistically; therefore, they were combined into one common control group. Statistical analyses were performed on the raw data before normalization. *,**,*** p<0.05, 0.01 and 0.001, compared to control. #, ##,### p<0.05, 0.01, and 0.001, respectively, compared to the LPS-treated group, n=3-8. Statistical analyses were conducted using one-way ANOVA followed by the Bonferroni post hoc test.

Our previous studies have shown that the hippocampus is highly sensitive to peripheral proinflammatory signaling (Czapski et al., 2013; Czapski et al., 2010; Czapski et al., 2016; Czapski et al., 2024; Czapski et al., 2020). Therefore, we analyzed whether LPS-induced systemic inflammatory response induces alterations in this structure. First, we measured the levels of BET proteins in the hippocampi of animals treated with LPS using commercial ELISA assay kits. Although our previous paper (Czapski et al., 2024) reported a significant increase in mRNA levels for *Brd4* 3 hours after LPS injection, the current study did not show a corresponding change in Brd4 levels at this early time point (data not shown). However, a significant increase in Brd4 protein levels was observed 12 h after LPS treatment (Fig. 3).

**Figure 3:**
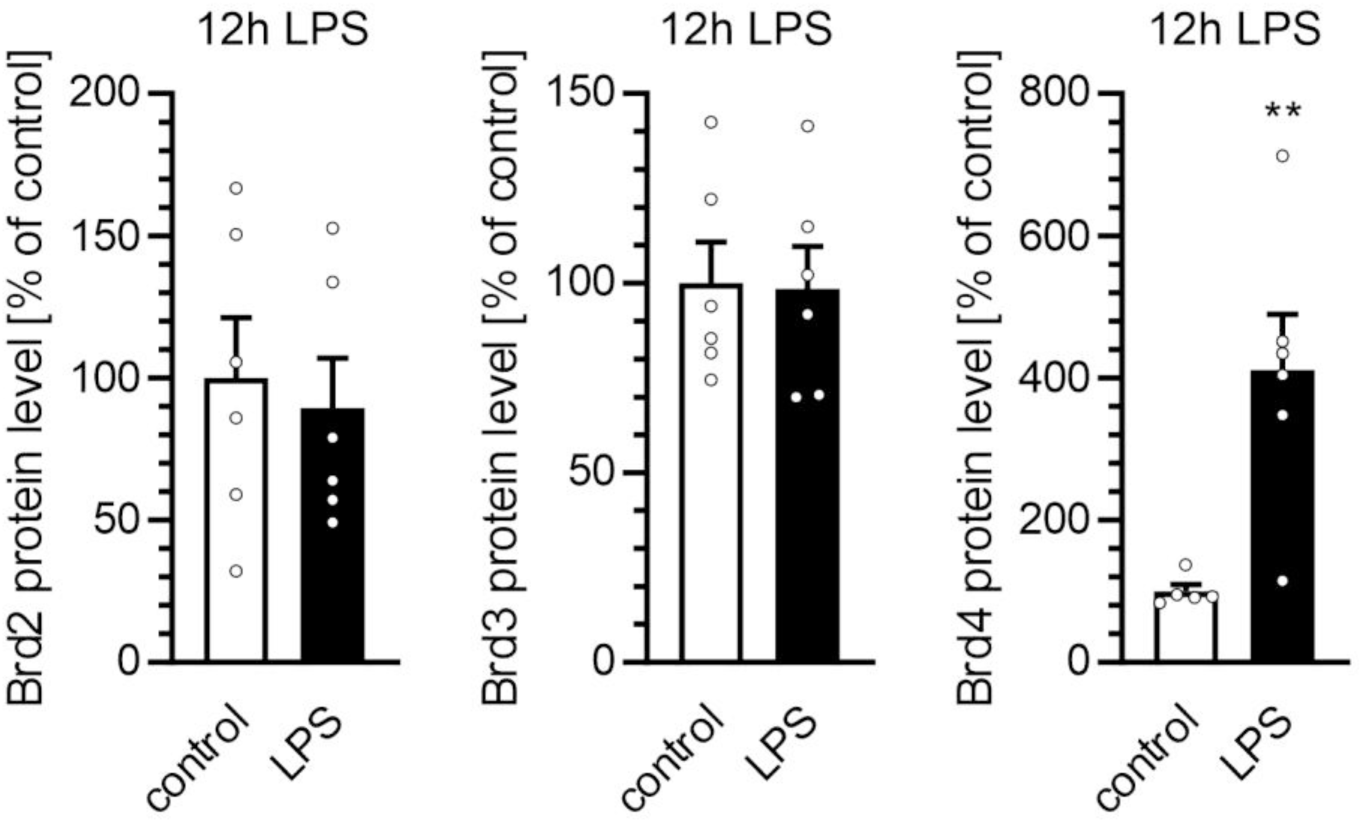
The effect of LPS-evoked systemic inflammation on the level of BET proteins in the murine hippocampus. Mice were injected with LPS (1 mg/kg body weight) or the respective volumes of vehicle. The levels of BET proteins in the hippocampus were determined 12 h post-injection using ELISA and normalized to the total protein level. ** p<0.01, compared to control, n=5-6. Statistical analyses were conducted using the Student’s t-test (Brd2, Brd3) or Mann-Whitney test (Brd4), selected based on data distribution.

Next, we analyzed the impact of LPS and JQ1 on processes related to inflammatory signaling in the hippocampus. Because the principal mode of action of BET inhibitors is through the modulation of gene expression, we focused on the quantitative analysis of the mRNA levels of selected genes, which seemed more reliable than measuring protein levels. We limited our analysis to the canonical markers of cytotoxic/cytoprotective states, *Il1b*, *Il6*, *Tnf*, *Nos2*, and *Arg1*. Similar to the upregulation of cytokines in the blood serum, the transcription of selected inflammation-related genes was also elevated during the systemic inflammatory response at both 3 and 12 h after LPS administration (Fig. 4a and 4b, respectively). The increased expression of these genes suggests that microglia are activated toward the proinflammatory/cytotoxic (often referred to as ‘M1’) phenotype. Simultaneously, the expression of Arg1, a marker of the microglia’s anti-inflammatory/cytoprotective phenotype (often referred to as ‘M2’), remained unchanged in the systemic inflammatory response group. We observed an evident inhibitory effect of JQ1 on the expression of proinflammatory genes 3 h after LPS treatment. However, this effect was no longer apparent after 12 hours. JQ1 treatment alone did not affect the expression of any of the tested genes under the control conditions.

**Figure 4:**
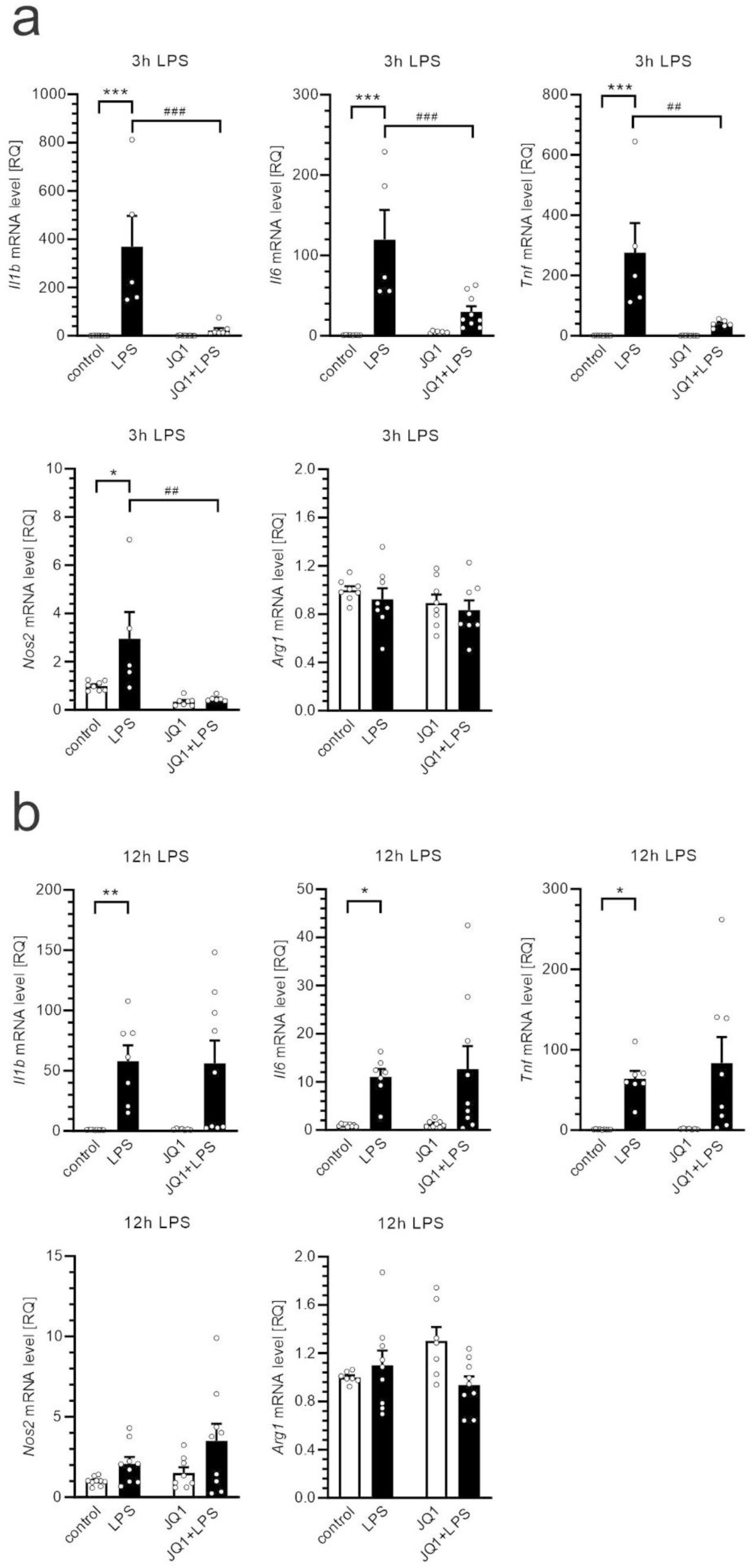
The effect of LPS and JQ1 on expression of inflammation-related genes in hippocampus. Mice were injected with JQ1 (50 mg/kg body weight) and after 30 min with LPS (1 mg/kg body weight). The respective groups received the respective vehicle volumes. The mRNA levels in the hippocampus were determined using qPCR at 3 h (a) and 12 h (b) after LPS administration. *,**,*** p<0.05, 0.01, and 0.001, comparing to respective control. ##, ### p<0.01, and 0.001, respectively, compared to the respective LPS-treated group, n=5-9 (a) and 7-9 (b). Statistical analyses were conducted using one-way ANOVA followed by the Bonferroni post hoc test. RQ, relative quantification.

The sickness behavior index (MSS) and inflammatory gene expression data from the animals in all experimental groups were subjected to correlation analysis. As expected, MSS was positively correlated with the mRNA levels of *Il1b*, *Il6*, and *Tnf*; however, the correlation was moderate, likely due to the relatively low number of data pairs (Supplementary Fig. 3).

As shown previously, using the same dose of LPS, the neuroinflammatory processes in the brain activated by the systemic inflammatory response significantly increase the phosphorylation of Tau protein at Ser396 (Czapski et al., 2016). In the present study, we confirmed and extended this observation. As presented in Fig. 5, 12 h after peripheral administration of LPS, the level of Tau protein in the hippocampus was not altered, but phosphorylation of Ser 396 and Ser 416 was significantly increased, whereas phosphorylation of Ser 199-202 demonstrated a strong tendency to increase (p=0.05). Pretreatment with JQ1 alone did not affect Tau levels or phosphorylation and had a negligible effect on the phosphorylation of this protein induced by a systemic inflammatory response.

**Figure 5:**
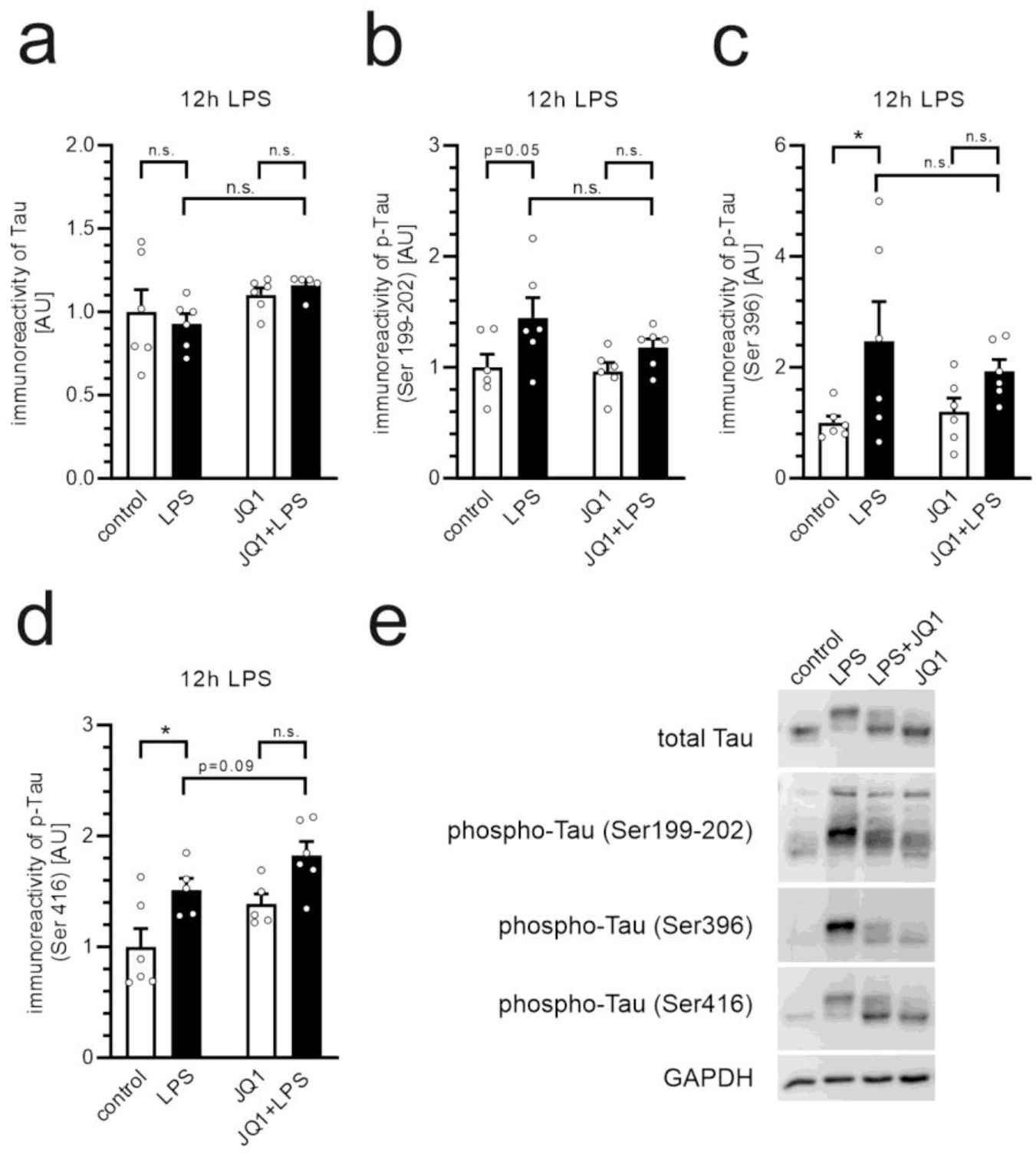
The effect of JQ1 on Tau level and phosphorylation in hippocampus after LPS-evoked systemic inflammation. Mice were injected with JQ1 (50 mg/kg body weight) and after 30 min with LPS (1 mg/kg body weight). The respective groups received the respective vehicle volumes. The level and phosphorylation of Tau in the hippocampus were determined using Western blotting with Gapdh as a loading control. The immunoreactivity of Tau (a) was normalized to the Gapdh level, and the immunoreactivity of phospho-Tau (b-d) was normalized to the total Tau level. * p<0.05, Statistical analyses were conducted using one-way ANOVA followed by the Bonferroni post hoc test. n=5-6 (a,d) and n=6 (b,c). (e) Typical images were presented. n.s. – not significant.

However, JQ1 significantly reduced the levels of Aβ_1-40_ and Aβ_1-42_ in the hippocampus of wild-type mice (Fig. 6a, 6b). This effect was evident in both control and LPS-treated animals. Notably, peripheral administration of LPS did not affect Aβ levels in the hippocampus. The levels of Aβ_1-40_ and Aβ_1-42_ in the hippocampi of the animals in all experimental groups were subjected to correlation analysis. As shown in Fig. 6c, the levels of Aβ1-40 and Aβ1-42 were highly positively correlated, which may suggest a common mechanism of JQ1 action for these two forms of Aβ. Because ELISA quantification relies on a standard curve, the confidence in quantifying our samples could be limited due to the very low concentrations, which are close to the lowest point on the standard curve. To mitigate this limitation, each sample was analyzed in duplicate (two technical replicates). The strong correlation between Aβ_1-40_ and Aβ_1-42_ levels increased the confidence in the reliability of our results.

**Figure 6:**
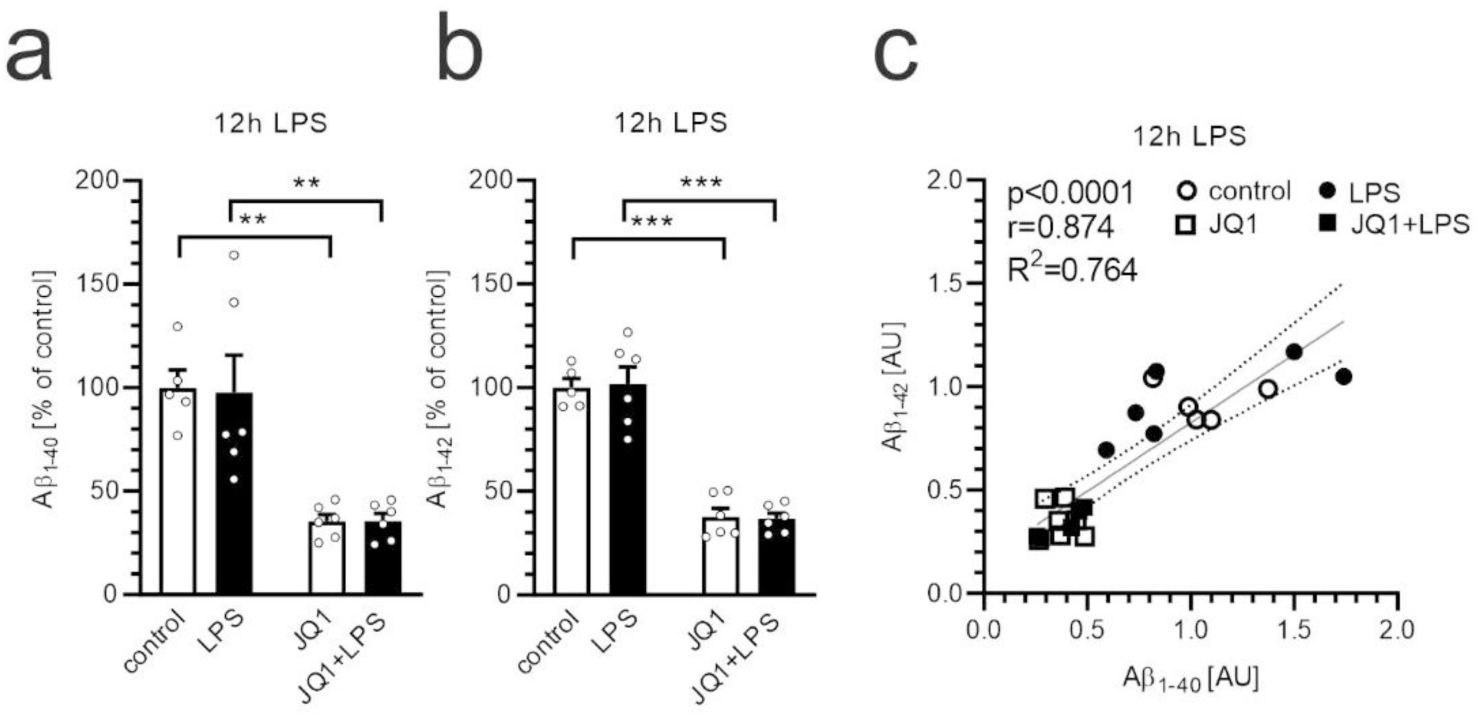
The effect of JQ1 on Amyloid-β level in hippocampus 12 h after LPS-evoked systemic inflammation. Mice were injected with JQ1 (50 mg/kg body weight) and after 30 min with LPS (1 mg/kg body weight). The respective groups received the respective vehicle volumes. The levels of Aβ_1-40_ (a) and Aβ_1-42_ (b) in the hippocampus were determined using ELISA. **,*** p<0.01 and 0.001, n=5-6. Statistical analyses were conducted using one-way ANOVA followed by the Bonferroni post hoc test. (c) Analysis of the correlation between Aβ_1-40_ and Aβ_1-42_ in the hippocampus. Dotted lines represent 95% confidence intervals.

Our previous study showed that JQ1 significantly attenuates phagocytic function in murine microglia in vitro (Matuszewska et al., 2022). We demonstrated that the phagocytosis of fluorescently labeled Aβ_1-42_ by BV2 cells was reduced after treatment with JQ1, and the expression of several phagocytosis/endocytosis-related genes was altered in BV2 cells exposed to JQ1. Moreover, our recent in vivo study, conducted under the same experimental conditions demonstrated that JQ1 reduced LPS-induced *Cd33* expression in the hippocampus but had no effect on the transcription of the *Cd33* gene in mice not exposed to endotoxemia (Czapski et al., 2024). Therefore, we hypothesized that alterations in phagocytosis- or endocytosis-related genes might cause JQ1-evoked reduction in Aβ load in the hippocampus. Thus, this study focused on investigating changes in the expression of genes previously implicated in the AD pathomechanism. As shown in Fig. 7, JQ1 did not affect the mRNA levels of any of the tested phagocytosis/endocytosis-related genes in the mouse hippocampus at either 3 h or 12 h after administration.

**Figure 7:**
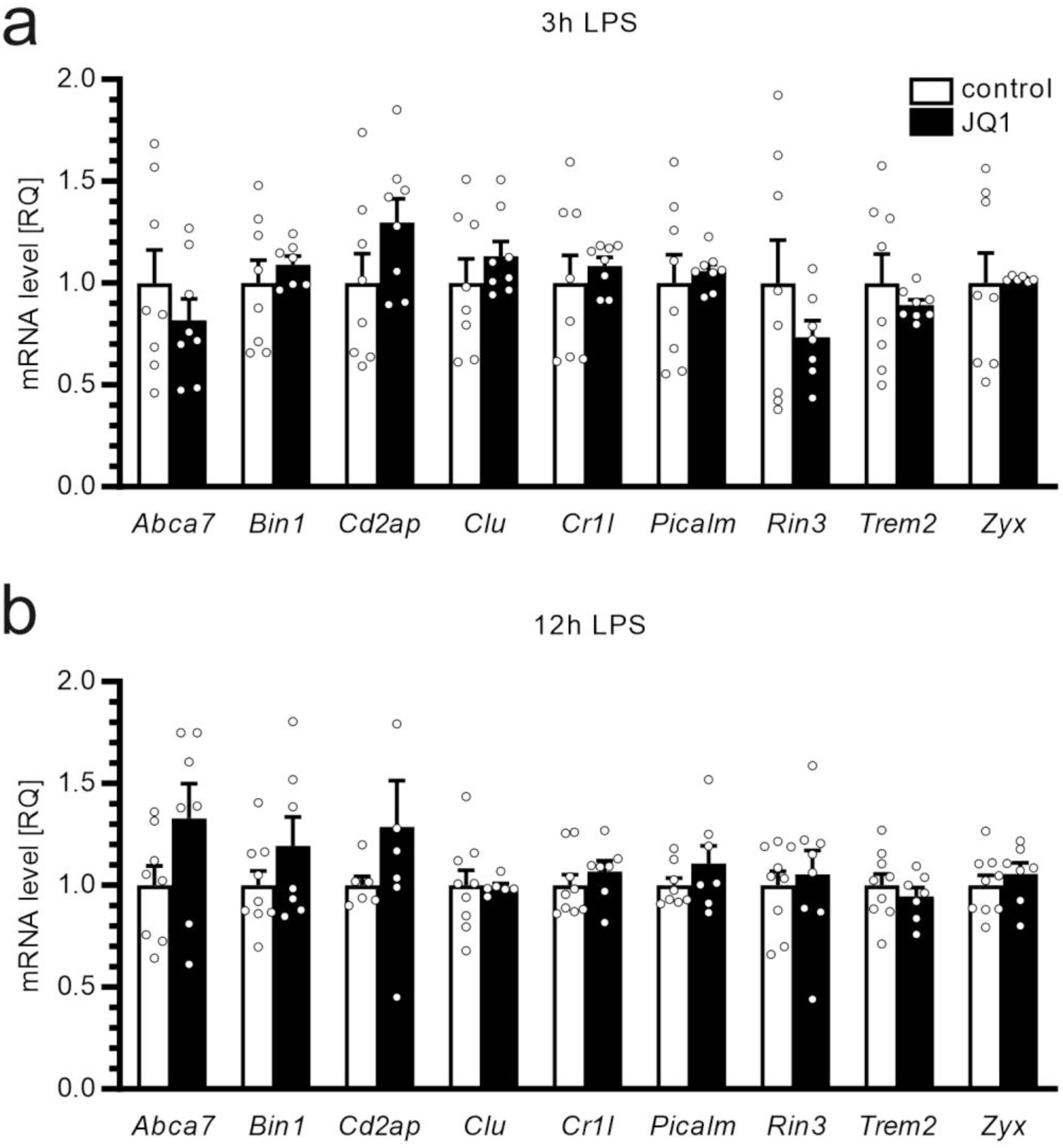
The effect of JQ1 on expression of phagocytosis-related genes in hippocampus. Mice were injected with JQ1 (50 mg/kg body weight) and NaCl after 30 min, as described above. The control group received the respective vehicle volume. Hippocampal mRNA levels were determined at 3 h (a) and 12 h (b) post-injection using qPCR. n=6-8 (a) and 6-9 (b). Statistical analyses were conducted using one-way ANOVA followed by the Bonferroni post hoc test. RQ, relative quantification.

Additionally, since the impact on amyloidogenesis is a potential mechanism of JQ1 action, we investigated the protein levels of β-secretase (BACE1) and γ-secretase (Aph1, Presenilin 1 and 2, PEN2, Nicastrin). However, we did not find any impact of LPS or JQ1 on the expression of those proteins (data not shown).

## Discussion

Activation of innate immune mechanisms is a component of virtually all neurodegenerative conditions (Castro-Gomez and Heneka, 2024). Pattern recognition receptor-mediated inflammatory responses are critical during both prodromal and clinical phases of these diseases. Recent data have highlighted the importance of the bacterial endotoxin, LPS, as a factor that may significantly contribute to or even promote neurodegeneration. The proposed ‘endotoxin hypothesis’ of AD suggests that LPS may be a crucial factor in triggering the pathological cascade leading to neurodegenerative injury (Brown, 2019; Brown and Heneka, 2024). Sepsis-associated encephalopathy, which occurs in approximately 70% of patients with severe sepsis, is related to long-lasting brain dysfunction, including cognitive deficits and post-traumatic stress disorder with increased anxiety (Grünewald et al., 2024; Iwashyna et al., 2010; Sipilä et al., 2021). Fine-tuning of inflammatory signaling is increasingly being considered as a promising strategy to change the trajectory of neurodegeneration. Therefore, identification of potential therapeutic targets to attenuate LPS-induced brain dysfunction and cognitive changes is necessary to promote healthy aging. BET proteins, the key players in the epigenetic control of gene transcription, have been proposed for inflammation-targeted treatment (Clayton et al., 2024; Martella et al., 2023).

Our study employed a mouse model of LPS-induced mild systemic inflammatory response, based on previous studies, in which we demonstrated that intraperitoneal injection of LPS (1.5×10^7^ U/kg body weight) evoked transient but robust and reproducible biochemical and molecular changes in the hippocampus of experimental animals (Czapski et al., 2013; Czapski et al., 2010; Czapski et al., 2016; Czapski et al., 2024; Czapski et al., 2020; Jacewicz et al., 2009). Moreover, we observed temporary symptoms of sickness behavior lasting up to 48 h and long-lasting cognitive impairments (Czapski et al., 2013; Czapski et al., 2007; Czapski et al., 2016; Czapski et al., 2020; Jacewicz et al., 2009). It is important to note that our model is an acute inflammatory response, not a model of the AD itself. However, BET inhibitors influenced some processes that are also relevant to AD. In this study, we used a small-molecule broad-spectrum inhibitor of BET proteins, JQ1. JQ1 exerts its effects by competitively binding to the acetyl-lysine recognition pocket of BET proteins, thereby preventing the binding of acetylated proteins and disrupting the recruitment of transcriptional complexes (Filippakopoulos et al., 2010). JQ1 was well tolerated without overt signs of toxicity or weight loss at a dose of 50 mg/kg in mice (Filippakopoulos et al., 2010). Pharmacokinetic analysis of JQ1 demonstrated its good permeability across the blood-brain barrier (AUC_brain_/AUC_plasma_ = 98%) after a single intraperitoneal dose of 50 mg/kg in male mice (Matzuk et al., 2012). However, it cannot be confidently excluded that some systemic interactions account for the effects observed in the brain.

The response of an organism to infection is complex and multiphase. A component of this response is a phenomenon known as sickness behavior (Dantzer, 2023), a universal and non-specific pattern of symptoms that develops during activation of the immune system. Fever, sleepiness, lethargy, anorexia, reduced social interest, etc. - all these symptoms are related to suppressed animal activity. They are considered to promote the fight against infection (energy repartition, metabolic reprogramming (O’Neill et al., 2016)) and prevent the spread of infectious agents in the population (Dantzer, 2023). The mechanism of sickness behavior in response to infectious agents is related to the immune system-dependent release of proinflammatory mediators, especially interleukin-1β (Dantzer et al., 1998), which affects brain function. Further research is required to gain a detailed understanding of the molecular mechanisms involved in this phenomenon. Our study investigated the role of BET proteins in sickness behavior, recognizing their crucial role in regulating numerous genes and pathways associated with immunity (Wang et al., 2021). Numerous studies have demonstrated that BET inhibitors block excessive production of inflammatory mediators and reduce the cell surface expression of chemokine receptors (Maksylewicz et al., 2019; Sanchez-Ventura et al., 2019; Wasiak et al., 2023). Following these studies, our results demonstrated that JQ1 significantly reduces LPS-evoked sickness behavior in mice, which is directly related to its anti-inflammatory effect, as shown by correlation analysis. However, we observed that JQ1 treatment affected only a select group of proinflammatory genes in a time-dependent manner. Intraperitoneal administration of JQ1 decreased serum levels and hippocampal gene expression of proinflammatory mediators in the LPS-treated group but did not affect the basal levels of these cytokines. We expect that a more powerful analysis (on larger experimental groups) or more extensive analysis (at more time points) could confirm this effect. Moreover, it would be reasonable to analyze the impact of JQ1 on long-lasting changes, such as LPS-evoked cognitive impairment, in future studies. The mechanisms underlying the effect of JQ1 on sickness behavior likely involve a reduction in cytokine levels but could also include other processes. These effects may include a decrease in vascular inflammation, modulation of the complement cascade, and normalization of cerebral blood flow (Huang et al., 2017; Wasiak et al., 2017; Yang et al., 2018).

The mechanism underlying the anti-inflammatory action of JQ1 is related to its effect on gene expression. Because BET proteins interact with a broad spectrum of transcription and elongation factors, their inhibition may affect various molecular pathways. Previous studies have shown that JQ1 suppresses LPS-induced microglial activation via the MAPK/NF-κB signaling pathway (Wang et al., 2018a). Activation of NF-κB and subsequent expression of inflammatory genes require acetylation of the RelA subunit at lysine-310 (Chen et al., 2002).

A notable finding of the current study was that a single peripheral administration of JQ1 decreased hippocampal levels of native murine Aβ in wild-type mice. This effect was evident for Aβ_1-40_ and Aβ_1-42_ both under control conditions and in LPS-treated animals. In the extraction protocol, we used a guanidine hydrochloride-containing buffer, assuming that we could measure the total level of Aβ in both the soluble and insoluble forms in the tissue. The amount of endogenous Aβ in wild-type mice is significantly smaller than that in transgenic animals overexpressing human Aβ; therefore, to assess Aβ levels, we used an ELISA assay, which has been previously demonstrated to be a specific and accurate method for measuring low levels of endogenous Aβ in wild-type mice (Lu et al., 2016; Peng et al., 2021). Although rodents do not naturally develop AD, an increase in Aβ levels is typically observed in aged wild-type animals (Drummond and Wisniewski, 2017; Fukumoto et al., 2004; Hoh Kam et al., 2010; Krstic et al., 2012; Oakley et al., 2006). However, with the exception of a single study by Ahlemeyer et al. (Ahlemeyer et al., 2018), extracellular deposits of Aβ have not been observed in the brains of naïve wild-type rodents. Immunological challenges—whether single prenatal or combined prenatal and postnatal (at 12 months)—have been shown to induce the formation of plaque-like structures in 15-month-old wild-type mice. However, the density of these “plaques” remained significantly lower compared to those observed in transgenic Alzheimer’s disease model mice (Krstic et al., 2012). Also, administering LPS to rodents increased Aβ levels in the brain; however, multiple injections of LPS were necessary (Hossain et al., 2018; Ifuku et al., 2012; Kahn et al., 2012; Lee et al., 2008). In contrast, a single, but very high dose of LPS increased soluble Aβ levels in the rat brain (Wang et al., 2018b). In our study, the levels of Aβ_1-40_ and Aβ_1-42_ did not change 12 h after peripheral administration of LPS. However, regardless of LPS-induced systemic inflammation, the administration of JQ1 significantly decreased Aβ levels in the hippocampus. Our previous studies have demonstrated that JQ1 significantly reduces *Cd33* gene expression in the microglial BV2 cell line (Matuszewska et al., 2022) and in mouse brains (Czapski et al., 2024). This suggests that the observed effect of JQ1 may involve Cd33, potentially linked to the activation of Aβ scavenging via microglial phagocytosis. Therefore, this study focused on other phagocytosis/endocytosis-related genes previously implicated in the AD pathomechanism (Jansen et al., 2019). First, we performed a curated analysis of the expression data deposited in NCBI’s Gene Expression Omnibus (Edgar et al., 2002) and accessible through the GEO Series accession number GSE155408 (https://www.ncbi.nlm.nih.gov/geo/query/acc.cgi?acc=GSE155408) (Table 1). The authors of that dataset, Baek and co-workers, used RNA sequencing to analyze the impact of JQ1 on gene expression in the human microglial HMC3 line (Baek et al., 2021). For comparison, we have included a summary of our previously published analysis of murine microglia (Matuszewska et al., 2022). The effect of JQ1 varied between the two cell lines. We hypothesize that these differences may be attributed to the effects of varying JQ1 doses and incubation times, as well as the testing of different organisms. Therefore, a mechanistic explanation for the anti-Aβ activity of JQ1 in the hippocampus requires further research.

**Table 1.**
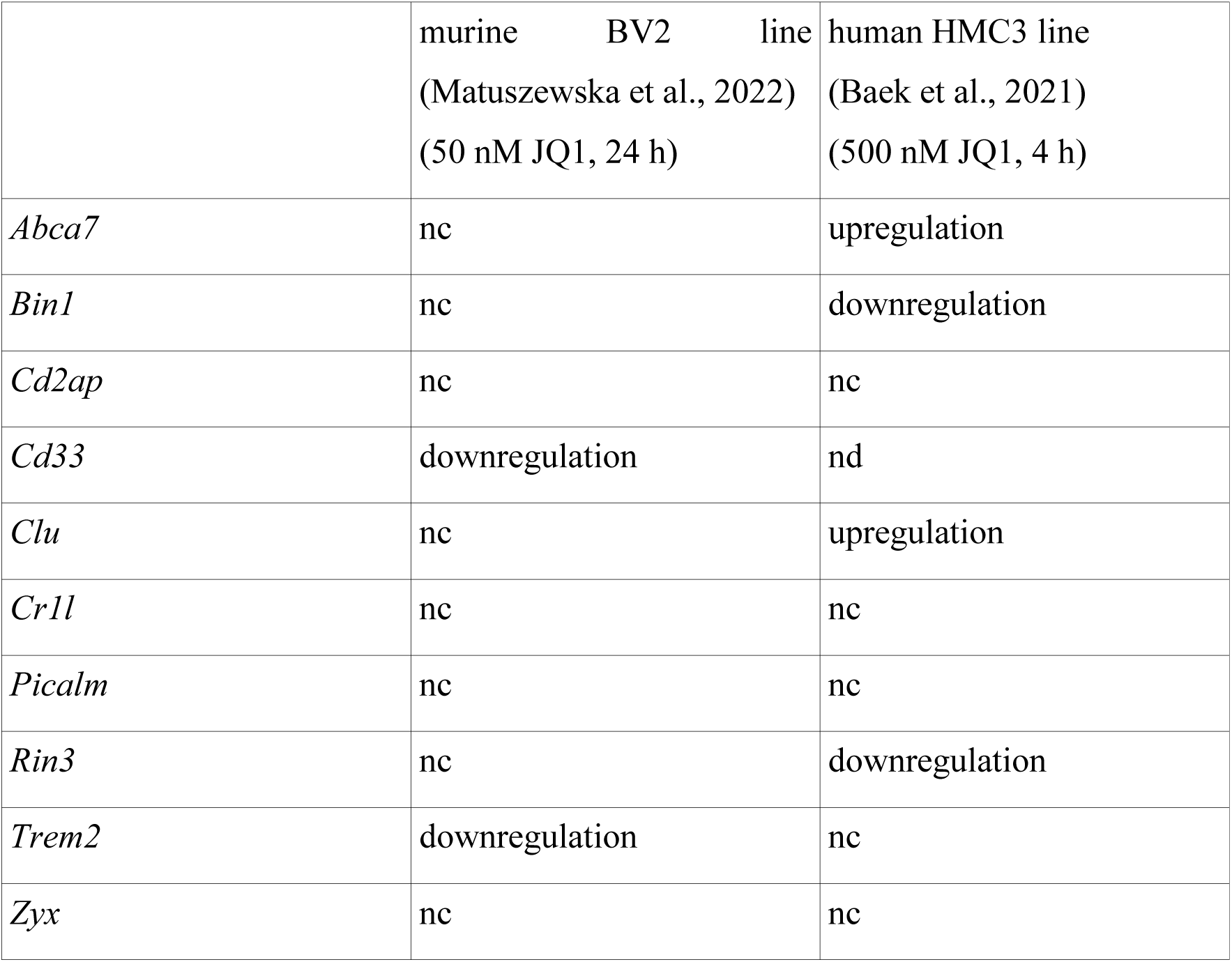
The curated analysis of the effect of JQ1 on the expression of phagocytosis-related genes in BV2 and HMC3 lines The curated analysis of the RNA sequencing data by Baek and co-workers (Baek et al., 2021) that have been deposited in NCBI’s GEO database and are accessible through GEO Series accession number GSE155408 (https://www.ncbi.nlm.nih.gov/geo/query/acc.cgi?acc=GSE155408). A summary of the qPCR data by Matuszewska and co-workers (Matuszewska et al., 2022) was added for comparison purposes. nc, not changed; nd, not detected. Details presented in Supplementary Table 3.

We propose that two other pathways may be involved in the anti-amyloid activity of JQ1. JQ1 might exert anti-amyloid effects by activating autophagy. Previous studies have demonstrated that JQ1 can stimulate autophagy by modulating the AMPK-mTOR-ULK1 signaling pathway (Li et al., 2020). Notably, dysregulation of autophagy has been implicated in the pathogenesis of AD (Di Meco et al., 2020). Consequently, activation of autophagy has emerged as a promising therapeutic approach for removing and clearing Aβ deposits (Friedman et al., 2015; Rahman et al., 2020). Indeed, numerous autophagy activators have been shown to effectively reduce Aβ levels in various experimental models (Kong et al., 2020; Ordóñez-Gutiérrez et al., 2018; Pierzynowska et al., 2019). A second potential pathway involves the inhibition of the NF-κB transcription factor, a key player in the pathogenesis of AD (Sun et al., 2022a). Notably, BET protein inhibitors attenuate the expression of NF-κB-regulated genes, thereby contributing to their anti-inflammatory effects. Furthermore, NF-κB inhibition has been shown to reduce Aβ production in cells overexpressing APP (Paris et al., 2007). Several NF-κB inhibitors have been demonstrated to decrease the levels of Aβ_1-40_, Aβ_1-42_, sAPPβ, and APP-CTFβ, suggesting that they diminish β-secretase-mediated cleavage of APP. It is noteworthy that the promoter of the gene encoding BACE1, the aspartyl protease responsible for β-site cleavage of APP and subsequent Aβ production, contains a binding site for NF-κB (Rossner et al., 2006). However, our experimental model did not show any increase in hippocampal Aβ levels during inflammation, suggesting that JQ1-mediated inhibition of inflammation-induced NF-κB and subsequent BACE1-dependent Aβ production is unlikely.

The anti-amyloidogenic effect of JQ1 in our experimental model appears to be specific to BET proteins, as preliminary data from another model (currently under preparation for publication) indicate that OTX-015, another BET inhibitor, exhibits a similar attenuating effect on Aβ levels in the brains of wild-type mice. However, BET inhibitors do not affect Tau phosphorylation in the hippocampus. Our previous study demonstrated that systemic inflammation enhances Tau phosphorylation at Se 396 by cyclin-dependent kinase 5 (Czapski et al., 2016). In the current study, we confirmed and extended this observation by demonstrating increased phosphorylation of Tau at Ser 199-202, Ser 396, and Ser 416 in the hippocampus during systemic inflammation. However, JQ1 did not affect Tau phosphorylation at any of the sites tested. The opposite effects of JQ1 were observed after chronic treatment of 3×Tg mice (Magistri et al., 2016). In the triple transgenic AD model, due to the presence of three human transgenes carrying mutations, *APP*(K670N/M671L; Swedish mutation), *PS1*(M146V), and *MAPT*(P301L), mice developed Aβ and Tau pathology followed by cognitive deficits. Mice treated with JQ1 for 15 weeks showed reduced Tau hyperphosphorylation at Ser 396 in the hippocampus and frontal cortex; however, JQ1 did not affect the level of soluble Aβ (Magistri et al., 2016). We propose that distinct molecular pathways may be responsible for Tau hyperphosphorylation in the transgenic 3×Tg and LPS-challenged mice.

Finally, our results demonstrated that only Brd4 is upregulated in the hippocampus under inflammatory conditions. This result is in agreement with the data of Wang et al. (Wang et al., 2020) that showed the selective upregulation of Brd4 in macrophages. Importantly, Brd4 plays a crucial role in controlling innate immune response to LPS (Bao et al., 2017). Compared to wild-type mice, Brd4-KO mice challenged with LPS displayed reduced expression of proinflammatory cytokine genes and were resistant to LPS-induced sepsis. However, selective gene silencing revealed that all three BET proteins regulate inflammatory gene expression in murine macrophages (Belkina et al., 2013). However, the action of JQ1 is not necessarily limited to Brd4, as Brd2 and Brd3—although not upregulated—may still play a role in regulating gene expression. Consequently, their inhibition might also influence JQ1’s effect in our experimental conditions. It is important to note that BET proteins have overlapping, yet distinct, roles in regulating gene expression. The differences arise from their specific domains and interaction partners. For example, our previous study demonstrated that silencing of the Brd2 and Brd4 genes in BV2 microglia reduced phagocytic activity to a greater extent than silencing of the Brd3 gene (Matuszewska et al., 2022). Moreover, the impact on phagocytosis-related gene expression was also specific to the BET protein. Because of the pivotal role of BET proteins in gene expression, the knock-out of Brd2 and Brd4 genes is lethal (Gyuris et al., 2009; Houzelstein et al., 2002; Shang et al., 2009), so the complex analysis of the role of particular BET proteins in vivo is currently not possible.

The limitation of this study is that it used a mouse model of systemic inflammation to explore aspects of human diseases. As previously suggested, inflammatory responses in mouse models show a limited correlation with human conditions, and the underlying regulatory mechanisms may differ significantly (Diehl and Boyle, 2018; Seok et al., 2013). However, other studies have indicated that some genomic responses during an inflammatory challenge show significant similarities between humans and mice, even though distinct regulatory mechanisms may be involved in the conserved transcriptional responses (Shay et al., 2013; Takao and Miyakawa, 2015). Moreover, although our results enhance the understanding of acute and dynamic changes in Aβ levels and their pharmacological regulators, the use of young mice limits the extrapolation of our findings to the chronic processes involved in Alzheimer’s disease. Therefore, further long-term studies are needed to assess their clinical relevance. Furthermore, although our attempts to elucidate the mechanism by which JQ1 reduces Aβ levels have been inconclusive, we have ruled out the impact on β- and γ-secretases, as well as the inhibition of NF-κB or phagocytosis. Future studies should explore other potential mechanisms, such as autophagy. Another limitation of our study is that we only tested male animals to minimize the variability associated with the estrous cycle in females. However, as AD is more prevalent in females, future studies should investigate both sexes to obtain more comprehensive results. Additionally, our study focused on acute changes, limiting our analysis to the short-term effects of JQ1 on animal behavior. Given that we previously observed cognitive impairment in mice persisting up to 7 days after LPS administration in this same model, future studies should investigate whether BET inhibition can mitigate these long-term effects.

## Conclusion

Our results indicate that inhibiting BET proteins is an effective strategy for attenuating the inflammatory response in both the periphery and the brain, including the alleviation of sickness behavior. Moreover, the inhibitor of BET proteins, JQ1, has been demonstrated to decrease the brain level of Aβ, suggesting that inhibition of BET proteins may be considered a potential anti-inflammatory and anti-amyloidogenic treatment. Inhibitors of BET proteins affect the expression of numerous genes, and their long-term effects may be extensive. Nevertheless, their widespread use in clinical trials for cancer highlights the potential of BET proteins as promising therapeutic targets.

## Ethics approval

All experiments conducted on animals were approved by the II Local Ethics Committee for Animal Experimentation in Warsaw (permission WAW2/060/2020), and were carried out following the EU Directive 2010/63/EU for animal experiments, and complied with ARRIVE (Animal Research: Reporting of In Vivo Experiments) guidelines. All efforts were made to minimize animal suffering and to reduce the number of animals.

## Consent for publication

Not applicable.

## Data Availability

The data supporting the findings of this study are available in the RepOD repository at https://repod.icm.edu.pl/dataset.xhtml?persistentId=doi:10.18150/I3B5JT

## Acknowledgments

Not applicable.

## Glossary

Ab: antibody
AD: Alzheimer’s disease
Aβ: amyloid-beta
BCA: bicinchoninic acid
BET: bromodomain and extraterminal domain proteins
DMSO: dimethyl sulfoxide
EOAD: early onset Alzheimer’s disease
LOAD: late-onset Alzheimer’s disease
LPS: lipopolysaccharide
MSS: murine sepsis score
nc: not changed
nd: not detected
NFT: neurofibrillary tangles
NSAID: non-steroidal anti-inflammatory drugs
RIPA: radioimmunoprecipitation assay
SP: sickness behavior

## References

Ahlemeyer, B., Halupczok, S., Rodenberg-Frank, E., Valerius, K.P., Baumgart-Vogt, E., 2018. Endogenous Murine Amyloid-β Peptide Assembles into Aggregates in the Aged C57BL/6J Mouse Suggesting These Animals as a Model to Study Pathogenesis of Amyloid-β Plaque Formation. J. Alzheimers Dis. 61, 1425–1450. doi: 10.3233/jad-170923.

Armstrong, R.A., 2014. A critical analysis of the ‘amyloid cascade hypothesis’. Folia Neuropathol. 52, 211–225. doi:

Back, M.R., Sarac, T.P., Moldawer, L.L., Welborn, M.B., 3rd, Seeger, J.M., Huber, T.S., 2000. Laparotomy prevents lethal endotoxemia in a murine sequential insult model by an IL-10-dependent mechanism. Shock 14, 157–162. doi: 10.1097/00024382-200014020-00014.

Baek, M., Yoo, E., Choi, H.I., An, G.Y., Chai, J.C., Lee, Y.S., Jung, K.H., Chai, Y.G., 2021. The BET inhibitor attenuates the inflammatory response and cell migration in human microglial HMC3 cell line. Sci. Rep. 11, 8828. doi: 10.1038/s41598-021-87828-1.

Bao, Y., Wu, X., Chen, J., Hu, X., Zeng, F., Cheng, J., Jin, H., Lin, X., Chen, L.F., 2017. Brd4 modulates the innate immune response through Mnk2-eIF4E pathway-dependent translational control of IκBα. Proc. Natl. Acad. Sci. U. S. A. 114, E3993–e4001. doi: 10.1073/pnas.1700109114.

Barroeta-Espar, I., Weinstock, L.D., Perez-Nievas, B.G., Meltzer, A.C., Siao Tick Chong, M., Amaral, A.C., Murray, M.E., Moulder, K.L., Morris, J.C., Cairns, N.J., Parisi, J.E., Lowe, V.J., Petersen, R.C., Kofler, J., Ikonomovic, M.D., López, O., Klunk, W.E., Mayeux, R.P., Frosch, M.P., Wood, L.B., Gomez-Isla, T., 2019. Distinct cytokine profiles in human brains resilient to Alzheimer’s pathology. Neurobiol. Dis. 121, 327–337. doi: 10.1016/j.nbd.2018.10.009.

Belkina, A.C., Nikolajczyk, B.S., Denis, G.V., 2013. BET protein function is required for inflammation: Brd2 genetic disruption and BET inhibitor JQ1 impair mouse macrophage inflammatory responses. J. Immunol. 190, 3670–3678. doi: 10.4049/jimmunol.1202838.

Brown, G.C., 2019. The endotoxin hypothesis of neurodegeneration. J. Neuroinflammation 16, 180. doi: 10.1186/s12974-019-1564-7.

Brown, G.C., Heneka, M.T., 2024. The endotoxin hypothesis of Alzheimer’s disease. Mol. Neurodegener. 19, 30. doi: 10.1186/s13024-024-00722-y.

Bu, X.L., Yao, X.Q., Jiao, S.S., Zeng, F., Liu, Y.H., Xiang, Y., Liang, C.R., Wang, Q.H., Wang, X., Cao, H.Y., Yi, X., Deng, B., Liu, C.H., Xu, J., Zhang, L.L., Gao, C.Y., Xu, Z.Q., Zhang, M., Wang, L., Tan, X.L., Xu, X., Zhou, H.D., Wang, Y.J., 2015. A study on the association between infectious burden and Alzheimer’s disease. Eur. J. Neurol. 22, 1519–1525. doi: 10.1111/ene.12477.

Castro-Gomez, S., Heneka, M.T., 2024. Innate immune activation in neurodegenerative diseases. Immunity 57, 790–814. doi: 10.1016/j.immuni.2024.03.010.

Chartier-Harlin, M.C., Crawford, F., Houlden, H., Warren, A., Hughes, D., Fidani, L., Goate, A., Rossor, M., Roques, P., Hardy, J., et al., 1991. Early-onset Alzheimer’s disease caused by mutations at codon 717 of the beta-amyloid precursor protein gene. Nature 353, 844–846. doi: 10.1038/353844a0.

Chen, L.F., Mu, Y., Greene, W.C., 2002. Acetylation of RelA at discrete sites regulates distinct nuclear functions of NF-kappaB. EMBO J. 21, 6539–6548. doi: 10.1093/emboj/cdf660.

Clayton, N., Pellei, D., Lin, Z., 2024. Histone acetylation, BET proteins, and periodontal inflammation. Mol Oral Microbiol 39, 180–189. doi: 10.1111/omi.12438.

Cunningham, C., Campion, S., Lunnon, K., Murray, C.L., Woods, J.F., Deacon, R.M., Rawlins, J.N., Perry, V.H., 2009. Systemic inflammation induces acute behavioral and cognitive changes and accelerates neurodegenerative disease. Biol Psychiatry 65, 304–312. doi: 10.1016/j.biopsych.2008.07.024.

Czapski, G.A., Adamczyk, A., Strosznajder, R.P., Strosznajder, J.B., 2013. Expression and activity of PARP family members in the hippocampus during systemic inflammation: their role in the regulation of prooxidative genes. Neurochem. Int. 62, 664–673. doi: 10.1016/j.neuint.2013.01.020.

Czapski, G.A., Cakala, M., Chalimoniuk, M., Gajkowska, B., Strosznajder, J.B., 2007. Role of nitric oxide in the brain during lipopolysaccharide-evoked systemic inflammation. J. Neurosci. Res. 85, 1694–1703. doi: 10.1002/jnr.21294.

Czapski, G.A., Cakala, M., Gajkowska, B., Strosznajder, J.B., 2006. Poly(ADP-ribose) polymerase-1 inhibition protects the brain against systemic inflammation. Neurochem. Int. 49, 751–755. doi: 10.1016/j.neuint.2006.06.006.

Czapski, G.A., Cakala, M., Kopczuk, D., Kaminska, M., Strosznajder, J.B., 2004. Inhibition of nitric oxide synthase prevents energy failure and oxidative damage evoked in the brain by lipopolysaccharide. Pol. J. Pharmacol. 56, 643–646. doi:

Czapski, G.A., Gajkowska, B., Strosznajder, J.B., 2010. Systemic administration of lipopolysaccharide induces molecular and morphological alterations in the hippocampus. Brain Res. 1356, 85–94. doi: 10.1016/j.brainres.2010.07.096.

Czapski, G.A., Gassowska, M., Wilkaniec, A., Chalimoniuk, M., Strosznajder, J.B., Adamczyk, A., 2016. The mechanisms regulating cyclin-dependent kinase 5 in hippocampus during systemic inflammatory response: The effect on inflammatory gene expression. Neurochem. Int. 93, 103–112. doi: 10.1016/j.neuint.2016.01.005.

Czapski, G.A., Matuszewska, M., Cieślik, M., Strosznajder, J.S., 2024. Inhibitor of bromodomain and extraterminal domain proteins decreases transcription of Cd33 in the brain of mice subjected to systemic inflammation; a promising strategy for neuroprotection. Folia Neuropathol. 62, 127–135. doi: 10.5114/fn.2024.138140.

Czapski, G.A., Zhao, Y., Lukiw, W.J., Strosznajder, J.B., 2020. Acute Systemic Inflammatory Response Alters Transcription Profile of Genes Related to Immune Response and Ca(2+) Homeostasis in Hippocampus; Relevance to Neurodegenerative Disorders. Int. J. Mol. Sci. 21. doi: 10.3390/ijms21217838.

Dantzer, R., 2023. Evolutionary Aspects of Infections: Inflammation and Sickness Behaviors. Curr. Top. Behav. Neurosci. 61, 1–14. doi: 10.1007/7854_2022_363.

Dantzer, R., Bluthe, R.M., Gheusi, G., Cremona, S., Laye, S., Parnet, P., Kelley, K.W., 1998. Molecular basis of sickness behavior. Ann. N. Y. Acad. Sci. 856, 132–138. doi:

Di Meco, A., Curtis, M.E., Lauretti, E., Praticò, D., 2020. Autophagy Dysfunction in Alzheimer’s Disease: Mechanistic Insights and New Therapeutic Opportunities. Biol Psychiatry 87, 797–807. doi: 10.1016/j.biopsych.2019.05.008.

Diehl, A.G., Boyle, A.P., 2018. Conserved and species-specific transcription factor co-binding patterns drive divergent gene regulation in human and mouse. Nucleic Acids Res. 46, 1878–1894. doi: 10.1093/nar/gky018.

Douros, A., Santella, C., Dell’Aniello, S., Azoulay, L., Renoux, C., Suissa, S., Brassard, P., 2021. Infectious Disease Burden and the Risk of Alzheimer’s Disease: A Population-Based Study. J. Alzheimers Dis. 81, 329–338. doi: 10.3233/jad-201534.

Drummond, E., Wisniewski, T., 2017. Alzheimer’s disease: experimental models and reality. Acta Neuropathol. 133, 155–175. doi: 10.1007/s00401-016-1662-x.

Edgar, R., Domrachev, M., Lash, A.E., 2002. Gene Expression Omnibus: NCBI gene expression and hybridization array data repository. Nucleic Acids Res. 30, 207–210. doi: 10.1093/nar/30.1.207.

Filippakopoulos, P., Qi, J., Picaud, S., Shen, Y., Smith, W.B., Fedorov, O., Morse, E.M., Keates, T., Hickman, T.T., Felletar, I., Philpott, M., Munro, S., McKeown, M.R., Wang, Y., Christie, A.L., West, N., Cameron, M.J., Schwartz, B., Heightman, T.D., La Thangue, N., French, C.A., Wiest, O., Kung, A.L., Knapp, S., Bradner, J.E., 2010. Selective inhibition of BET bromodomains. Nature 468, 1067–1073. doi: 10.1038/nature09504.

Friedman, L.G., Qureshi, Y.H., Yu, W.H., 2015. Promoting autophagic clearance: viable therapeutic targets in Alzheimer’s disease. Neurotherapeutics 12, 94–108. doi: 10.1007/s13311-014-0320-z.

Fukumoto, H., Rosene, D.L., Moss, M.B., Raju, S., Hyman, B.T., Irizarry, M.C., 2004. Beta-secretase activity increases with aging in human, monkey, and mouse brain. Am. J. Pathol. 164, 719–725. doi: 10.1016/s0002-9440(10)63159-8.

Gąssowska-Dobrowolska, M., Kolasa-Wołosiuk, A., Cieślik, M., Dominiak, A., Friedland, K., Adamczyk, A., 2021. Alterations in Tau Protein Level and Phosphorylation State in the Brain of the Autistic-Like Rats Induced by Prenatal Exposure to Valproic Acid. Int. J. Mol. Sci. 22. doi: 10.3390/ijms22063209.

Gilan, O., Rioja, I., Knezevic, K., Bell, M.J., Yeung, M.M., Harker, N.R., Lam, E.Y.N., Chung, C.W., Bamborough, P., Petretich, M., Urh, M., Atkinson, S.J., Bassil, A.K., Roberts, E.J., Vassiliadis, D., Burr, M.L., Preston, A.G.S., Wellaway, C., Werner, T., Gray, J.R., Michon, A.M., Gobbetti, T., Kumar, V., Soden, P.E., Haynes, A., Vappiani, J., Tough, D.F., Taylor, S., Dawson, S.J., Bantscheff, M., Lindon, M., Drewes, G., Demont, E.H., Daniels, D.L., Grandi, P., Prinjha, R.K., Dawson, M.A., 2020. Selective targeting of BD1 and BD2 of the BET proteins in cancer and immunoinflammation. Science 368, 387–394. doi: 10.1126/science.aaz8455.

Griciuc, A., Serrano-Pozo, A., Parrado, A.R., Lesinski, A.N., Asselin, C.N., Mullin, K., Hooli, B., Choi, S.H., Hyman, B.T., Tanzi, R.E., 2013. Alzheimer’s disease risk gene CD33 inhibits microglial uptake of amyloid beta. Neuron 78, 631–643. doi: 10.1016/j.neuron.2013.04.014.

Grünewald, B., Wickel, J., Hahn, N., Rahmati, V., Rupp, H., Chung, H.Y., Haselmann, H., Strauss, A.S., Schmidl, L., Hempel, N., Grünewald, L., Urbach, A., Bauer, M., Toyka, K.V., Blaess, M., Claus, R.A., König, R., Geis, C., 2024. Targeted rescue of synaptic plasticity improves cognitive decline in sepsis-associated encephalopathy. Mol. Ther. doi: 10.1016/j.ymthe.2024.05.001.

Gyuris, A., Donovan, D.J., Seymour, K.A., Lovasco, L.A., Smilowitz, N.R., Halperin, A.L., Klysik, J.E., Freiman, R.N., 2009. The chromatin-targeting protein Brd2 is required for neural tube closure and embryogenesis. Biochim. Biophys. Acta 1789, 413–421. doi: 10.1016/j.bbagrm.2009.03.005.

Harden, L.M., Kent, S., Pittman, Q.J., Roth, J., 2015. Fever and sickness behavior: Friend or foe? Brain Behav. Immun. 50, 322–333. doi: 10.1016/j.bbi.2015.07.012.

Hoh Kam, J., Lenassi, E., Jeffery, G., 2010. Viewing ageing eyes: diverse sites of amyloid Beta accumulation in the ageing mouse retina and the up-regulation of macrophages. PloS one 5. doi: 10.1371/journal.pone.0013127.

Holmes, C., Cunningham, C., Zotova, E., Woolford, J., Dean, C., Kerr, S., Culliford, D., Perry, V.H., 2009. Systemic inflammation and disease progression in Alzheimer disease. Neurology 73, 768–774. doi: 10.1212/WNL.0b013e3181b6bb95.

Hossain, M.S., Tajima, A., Kotoura, S., Katafuchi, T., 2018. Oral ingestion of plasmalogens can attenuate the LPS-induced memory loss and microglial activation. Biochem. Biophys. Res. Commun. 496, 1033–1039. doi: 10.1016/j.bbrc.2018.01.078.

Houzelstein, D., Bullock, S.L., Lynch, D.E., Grigorieva, E.F., Wilson, V.A., Beddington, R.S., 2002. Growth and early postimplantation defects in mice deficient for the bromodomain-containing protein Brd4. Mol. Cell. Biol. 22, 3794–3802. doi: 10.1128/mcb.22.11.3794-3802.2002.

Huang, M., Zeng, S., Zou, Y., Shi, M., Qiu, Q., Xiao, Y., Chen, G., Yang, X., Liang, L., Xu, H., 2017. The suppression of bromodomain and extra-terminal domain inhibits vascular inflammation by blocking NF-κB and MAPK activation. British journal of pharmacology 174, 101–115. doi: 10.1111/bph.13657.

Ifuku, M., Katafuchi, T., Mawatari, S., Noda, M., Miake, K., Sugiyama, M., Fujino, T., 2012. Anti-inflammatory/anti-amyloidogenic effects of plasmalogens in lipopolysaccharide-induced neuroinflammation in adult mice. J. Neuroinflammation 9, 197. doi: 10.1186/1742-2094-9-197.

Iwashyna, T.J., Ely, E.W., Smith, D.M., Langa, K.M., 2010. Long-term cognitive impairment and functional disability among survivors of severe sepsis. Jama 304, 1787–1794. doi: 10.1001/jama.2010.1553.

Jacewicz, M., Czapski, G.A., Katkowska, I., Strosznajder, R.P., 2009. Systemic administration of lipopolysaccharide impairs glutathione redox state and object recognition in male mice. The effect of PARP-1 inhibitor. Folia Neuropathol. 47, 321–328. doi:

Jansen, I.E., Savage, J.E., Watanabe, K., Bryois, J., Williams, D.M., Steinberg, S., Sealock, J., Karlsson, I.K., Hägg, S., Athanasiu, L., Voyle, N., Proitsi, P., Witoelar, A., Stringer, S., Aarsland, D., Almdahl, I.S., Andersen, F., Bergh, S., Bettella, F., Bjornsson, S., Brækhus, A., Bråthen, G., de Leeuw, C., Desikan, R.S., Djurovic, S., Dumitrescu, L., Fladby, T., Hohman, T.J., Jonsson, P.V., Kiddle, S.J., Rongve, A., Saltvedt, I., Sando, S.B., Selbæk, G., Shoai, M., Skene, N.G., Snaedal, J., Stordal, E., Ulstein, I.D., Wang, Y., White, L.R., Hardy, J., Hjerling-Leffler, J., Sullivan, P.F., van der Flier, W.M., Dobson, R., Davis, L.K., Stefansson, H., Stefansson, K., Pedersen, N.L., Ripke, S., Andreassen, O.A., Posthuma, D., 2019. Genome-wide meta-analysis identifies new loci and functional pathways influencing Alzheimer’s disease risk. Nat. Genet. 51, 404–413. doi: 10.1038/s41588-018-0311-9.

Jostes, S., Nettersheim, D., Fellermeyer, M., Schneider, S., Hafezi, F., Honecker, F., Schumacher, V., Geyer, M., Kristiansen, G., Schorle, H., 2017. The bromodomain inhibitor JQ1 triggers growth arrest and apoptosis in testicular germ cell tumours in vitro and in vivo. J. Cell. Mol. Med. 21, 1300–1314. doi: 10.1111/jcmm.13059.

Jucker, M., Walker, L.C., 2023. Alzheimer’s disease: From immunotherapy to immunoprevention. Cell 186, 4260–4270. doi: 10.1016/j.cell.2023.08.021.

Kahn, M.S., Kranjac, D., Alonzo, C.A., Haase, J.H., Cedillos, R.O., McLinden, K.A., Boehm, G.W., Chumley, M.J., 2012. Prolonged elevation in hippocampal Aβ and cognitive deficits following repeated endotoxin exposure in the mouse. Behav. Brain Res. 229, 176–184. doi: 10.1016/j.bbr.2012.01.010.

Kepp, K.P., Robakis, N.K., Høilund-Carlsen, P.F., Sensi, S.L., Vissel, B., 2023. The amyloid cascade hypothesis: an updated critical review. Brain 146, 3969–3990. doi: 10.1093/brain/awad159.

Kong, J., Wan, L., Wang, Y., Zhang, H., Zhang, W., 2020. Liraglutide Attenuates Aβ42 Generation in APPswe/SH-SY5Y Cells Through the Regulation of Autophagy. Neuropsychiatr Dis Treat 16, 1817–1825. doi: 10.2147/ndt.S260160.

Korb, E., Herre, M., Zucker-Scharff, I., Darnell, R.B., Allis, C.D., 2015. BET protein Brd4 activates transcription in neurons and BET inhibitor Jq1 blocks memory in mice. Nat. Neurosci. 18, 1464–1473. doi: 10.1038/nn.4095.

Krstic, D., Madhusudan, A., Doehner, J., Vogel, P., Notter, T., Imhof, C., Manalastas, A., Hilfiker, M., Pfister, S., Schwerdel, C., Riether, C., Meyer, U., Knuesel, I., 2012. Systemic immune challenges trigger and drive Alzheimer-like neuropathology in mice. J. Neuroinflammation 9, 151. doi: 10.1186/1742-2094-9-151.

Lee, J.W., Lee, Y.K., Yuk, D.Y., Choi, D.Y., Ban, S.B., Oh, K.W., Hong, J.T., 2008. Neuro-inflammation induced by lipopolysaccharide causes cognitive impairment through enhancement of beta-amyloid generation. J. Neuroinflammation 5, 37. doi: 10.1186/1742-2094-5-37.

Li, Q., Li, L., Fei, X., Zhang, Y., Qi, C., Hua, S., Gong, F., Fang, M., 2018. Inhibition of autophagy with 3-methyladenine is protective in a lethal model of murine endotoxemia and polymicrobial sepsis. Innate Immun 24, 231–239. doi: 10.1177/1753425918771170.

Li, Y., Xiang, J., Zhang, J., Lin, J., Wu, Y., Wang, X., 2020. Inhibition of Brd4 by JQ1 Promotes Functional Recovery From Spinal Cord Injury by Activating Autophagy. Frontiers in cellular neuroscience 14, 555591. doi: 10.3389/fncel.2020.555591.

Liu, L., Yang, C., Candelario-Jalil, E., 2021. Role of BET Proteins in Inflammation and CNS Diseases. Front Mol Biosci 8, 748449. doi: 10.3389/fmolb.2021.748449.

Lu, Y., Hoyte, K., Montgomery, W.H., Luk, W., He, D., Meilandt, W.J., Zuchero, Y.J., Atwal, J.K., Scearce-Levie, K., Watts, R.J., DeForge, L.E., 2016. Characterization of a sensitive mouse Aβ40 PD biomarker assay for Alzheimer’s disease drug development in wild-type mice. Bioanalysis 8, 1067–1075. doi: 10.4155/bio-2016-0003.

Magistri, M., Velmeshev, D., Makhmutova, M., Patel, P., Sartor, G.C., Volmar, C.H., Wahlestedt, C., Faghihi, M.A., 2016. The BET-Bromodomain Inhibitor JQ1 Reduces Inflammation and Tau Phosphorylation at Ser396 in the Brain of the 3xTg Model of Alzheimer’s Disease. Curr Alzheimer Res 13, 985–995. doi: 10.2174/1567205013666160427101832.

Maksylewicz, A., Bysiek, A., Lagosz, K.B., Macina, J.M., Kantorowicz, M., Bereta, G., Sochalska, M., Gawron, K., Chomyszyn-Gajewska, M., Potempa, J., Grabiec, A.M., 2019. BET Bromodomain Inhibitors Suppress Inflammatory Activation of Gingival Fibroblasts and Epithelial Cells From Periodontitis Patients. Front Immunol 10, 933. doi: 10.3389/fimmu.2019.00933.

Martella, N., Pensabene, D., Varone, M., Colardo, M., Petraroia, M., Sergio, W., La Rosa, P., Moreno, S., Segatto, M., 2023. Bromodomain and Extra-Terminal Proteins in Brain Physiology and Pathology: BET-ing on Epigenetic Regulation. Biomedicines 11. doi: 10.3390/biomedicines11030750.

Matuszewska, M., Cieślik, M., Wilkaniec, A., Strawski, M., Czapski, G.A., 2022. The Role of Bromodomain and Extraterminal (BET) Proteins in Controlling the Phagocytic Activity of Microglia In Vitro: Relevance to Alzheimer’s Disease. Int. J. Mol. Sci. 24. doi: 10.3390/ijms24010013.

Matzuk, M.M., McKeown, M.R., Filippakopoulos, P., Li, Q., Ma, L., Agno, J.E., Lemieux, M.E., Picaud, S., Yu, R.N., Qi, J., Knapp, S., Bradner, J.E., 2012. Small-molecule inhibition of BRDT for male contraception. Cell 150, 673–684. doi: 10.1016/j.cell.2012.06.045.

McGeer, E.G., McGeer, P.L., 1998. The importance of inflammatory mechanisms in Alzheimer disease. Exp. Gerontol. 33, 371–378. doi: 10.1016/s0531-5565(98)00013-8.

O’Neill, L.A., Kishton, R.J., Rathmell, J., 2016. A guide to immunometabolism for immunologists. Nat. Rev. Immunol. 16, 553–565. doi: 10.1038/nri.2016.70.

Oakley, H., Cole, S.L., Logan, S., Maus, E., Shao, P., Craft, J., Guillozet-Bongaarts, A., Ohno, M., Disterhoft, J., Van Eldik, L.J., Berr, y.R., Vassar, R., 2006. Intraneuronal beta-amyloid aggregates, neurodegeneration, and neuron loss in transgenic mice with five familial Alzheimer’s disease mutations: potential factors in amyloid plaque formation. J. Neurosci. 26, 10129–10140. doi:

Ordóñez-Gutiérrez, L., Benito-Cuesta, I., Abad, J.L., Casas, J., Fábrias, G., Wandosell, F., 2018. Dihydroceramide Desaturase 1 Inhibitors Reduce Amyloid-β Levels in Primary Neurons from an Alzheimer’s Disease Transgenic Model. Pharm. Res. 35, 49. doi: 10.1007/s11095-017-2312-2.

Paris, D., Patel, N., Quadros, A., Linan, M., Bakshi, P., Ait-Ghezala, G., Mullan, M., 2007. Inhibition of Abeta production by NF-kappaB inhibitors. Neurosci. Lett. 415, 11–16. doi: 10.1016/j.neulet.2006.12.029.

Peng, H.B., de Lannoy, I.A.M., Pang, K.S., 2021. Measuring Amyloid-β Peptide Concentrations in Murine Brain with Improved ELISA Assay. Curr Protoc 1, e253. doi: 10.1002/cpz1.253.

Perez-Nievas, B.G., Stein, T.D., Tai, H.C., Dols-Icardo, O., Scotton, T.C., Barroeta-Espar, I., Fernandez-Carballo, L., de Munain, E.L., Perez, J., Marquie, M., Serrano-Pozo, A., Frosch, M.P., Lowe, V., Parisi, J.E., Petersen, R.C., Ikonomovic, M.D., López, O.L., Klunk, W., Hyman, B.T., Gómez-Isla, T., 2013. Dissecting phenotypic traits linked to human resilience to Alzheimer’s pathology. Brain 136, 2510–2526. doi: 10.1093/brain/awt171.

Pierzynowska, K., Podlacha, M., Gaffke, L., Majkutewicz, I., Mantej, J., Węgrzyn, A., Osiadły, M., Myślińska, D., Węgrzyn, G., 2019. Autophagy-dependent mechanism of genistein-mediated elimination of behavioral and biochemical defects in the rat model of sporadic Alzheimer’s disease. Neuropharmacology 148, 332–346. doi: 10.1016/j.neuropharm.2019.01.030.

Qin, L., Wu, X., Block, M.L., Liu, Y., Breese, G.R., Hong, J.S., Knapp, D.J., Crews, F.T., 2007. Systemic LPS causes chronic neuroinflammation and progressive neurodegeneration. Glia 55, 453–462. doi: 10.1002/glia.20467.

Rahman, M.A., Rahman, M.S., Rahman, M.D.H., Rasheduzzaman, M., Mamun-Or-Rashid, A., Uddin, M.J., Rahman, M.R., Hwang, H., Pang, M.G., Rhim, H., 2020. Modulatory Effects of Autophagy on APP Processing as a Potential Treatment Target for Alzheimer’s Disease. Biomedicines 9. doi: 10.3390/biomedicines9010005.

Rossner, S., Sastre, M., Bourne, K., Lichtenthaler, S.F., 2006. Transcriptional and translational regulation of BACE1 expression--implications for Alzheimer’s disease. Prog. Neurobiol. 79, 95–111. doi: 10.1016/j.pneurobio.2006.06.001.

Sanchez-Ventura, J., Amo-Aparicio, J., Navarro, X., Penas, C., 2019. BET protein inhibition regulates cytokine production and promotes neuroprotection after spinal cord injury. J. Neuroinflammation 16, 124. doi: 10.1186/s12974-019-1511-7.

Seok, J., Warren, H.S., Cuenca, A.G., Mindrinos, M.N., Baker, H.V., Xu, W., Richards, D.R., McDonald-Smith, G.P., Gao, H., Hennessy, L., Finnerty, C.C., López, C.M., Honari, S., Moore, E.E., Minei, J.P., Cuschieri, J., Bankey, P.E., Johnson, J.L., Sperry, J., Nathens, A.B., Billiar, T.R., West, M.A., Jeschke, M.G., Klein, M.B., Gamelli, R.L., Gibran, N.S., Brownstein, B.H., Miller-Graziano, C., Calvano, S.E., Mason, P.H., Cobb, J.P., Rahme, L.G., Lowry, S.F., Maier, R.V., Moldawer, L.L., Herndon, D.N., Davis, R.W., Xiao, W., Tompkins, R.G., 2013. Genomic responses in mouse models poorly mimic human inflammatory diseases. Proc. Natl. Acad. Sci. U. S. A. 110, 3507–3512. doi: 10.1073/pnas.1222878110.

Shang, E., Wang, X., Wen, D., Greenberg, D.A., Wolgemuth, D.J., 2009. Double bromodomain-containing gene Brd2 is essential for embryonic development in mouse. Dev. Dyn. 238, 908–917. doi: 10.1002/dvdy.21911.

Shay, T., Jojic, V., Zuk, O., Rothamel, K., Puyraimond-Zemmour, D., Feng, T., Wakamatsu, E., Benoist, C., Koller, D., Regev, A., 2013. Conservation and divergence in the transcriptional programs of the human and mouse immune systems. Proc. Natl. Acad. Sci. U. S. A. 110, 2946–2951. doi: 10.1073/pnas.1222738110.

Sherrington, R., Rogaev, E.I., Liang, Y., Rogaeva, E.A., Levesque, G., Ikeda, M., Chi, H., Lin, C., Li, G., Holman, K., Tsuda, T., Mar, L., Foncin, J.F., Bruni, A.C., Montesi, M.P., Sorbi, S., Rainero, I., Pinessi, L., Nee, L., Chumakov, I., Pollen, D., Brookes, A., Sanseau, P., Polinsky, R.J., Wasco, W., Da Silva, H.A., Haines, J.L., Perkicak-Vance, M.A., Tanzi, R.E., Roses, A.D., Fraser, P.E., Rommens, J.M., St George-Hyslop, P.H., 1995. Cloning of a gene bearing missense mutations in early-onset familial Alzheimer’s disease. Nature 375, 754–760. doi: 10.1038/375754a0.

Shrum, B., Anantha, R.V., Xu, S.X., Donnelly, M., Haeryfar, S.M., McCormick, J.K., Mele, T., 2014. A robust scoring system to evaluate sepsis severity in an animal model. BMC Res. Notes 7, 233. doi: 10.1186/1756-0500-7-233.

Sipilä, P.N., Heikkilä, N., Lindbohm, J.V., Hakulinen, C., Vahtera, J., Elovainio, M., Suominen, S., Väänänen, A., Koskinen, A., Nyberg, S.T., Pentti, J., Strandberg, T.E., Kivimäki, M., 2021. Hospital-treated infectious diseases and the risk of dementia: a large, multicohort, observational study with a replication cohort. Lancet Infect. Dis. 21, 1557–1567. doi: 10.1016/s1473-3099(21)00144-4.

Sun, E., Motolani, A., Campos, L., Lu, T., 2022a. The Pivotal Role of NF-kB in the Pathogenesis and Therapeutics of Alzheimer’s Disease. Int. J. Mol. Sci. 23. doi: 10.3390/ijms23168972.

Sun, J., Ludvigsson, J.F., Ingre, C., Piehl, F., Wirdefeldt, K., Zagai, U., Ye, W., Fang, F., 2022b. Hospital-treated infections in early- and mid-life and risk of Alzheimer’s disease, Parkinson’s disease, and amyotrophic lateral sclerosis: A nationwide nested case-control study in Sweden. PLoS Med. 19, e1004092. doi: 10.1371/journal.pmed.1004092.

Takao, K., Miyakawa, T., 2015. Genomic responses in mouse models greatly mimic human inflammatory diseases. Proc. Natl. Acad. Sci. U. S. A. 112, 1167–1172. doi: 10.1073/pnas.1401965111.

Teeling, J.L., Perry, V.H., 2009. Systemic infection and inflammation in acute CNS injury and chronic neurodegeneration: underlying mechanisms. Neuroscience 158, 1062–1073. doi: 10.1016/j.neuroscience.2008.07.031.

Wang, H., Fu, H., Zhu, R., Wu, X., Ji, X., Li, X., Jiang, H., Lin, Z., Tang, X., Sun, S., Chen, J., Wang, X., Li, Q., Ji, Y., Chen, H., 2020. BRD4 contributes to LPS-induced macrophage senescence and promotes progression of atherosclerosis-associated lipid uptake. Aging 12, 9240–9259. doi: 10.18632/aging.103200.

Wang, H., Huang, W., Liang, M., Shi, Y., Zhang, C., Li, Q., Liu, M., Shou, Y., Yin, H., Zhu, X., Sun, X., Hu, Y., Shen, Z., 2018a. (+)-JQ1 attenuated LPS-induced microglial inflammation via MAPK/NFκB signaling. Cell Biosci 8, 60. doi: 10.1186/s13578-018-0258-7.

Wang, L.M., Wu, Q., Kirk, R.A., Horn, K.P., Ebada Salem, A.H., Hoffman, J.M., Yap, J.T., Sonnen, J.A., Towner, R.A., Bozza, F.A., Rodrigues, R.S., Morton, K.A., 2018b. Lipopolysaccharide endotoxemia induces amyloid-β and p-tau formation in the rat brain. Am. J. Nucl. Med. Mol. Imaging 8, 86–99. doi:

Wang, N., Wu, R., Tang, D., Kang, R., 2021. The BET family in immunity and disease. Signal Transduct Target Ther 6, 23. doi: 10.1038/s41392-020-00384-4.

Wang, Z.Q., Zhang, Z.C., Wu, Y.Y., Pi, Y.N., Lou, S.H., Liu, T.B., Lou, G., Yang, C., 2023. Bromodomain and extraterminal (BET) proteins: biological functions, diseases, and targeted therapy. Signal Transduct Target Ther 8, 420. doi: 10.1038/s41392-023-01647-6.

Wasiak, S., Fu, L., Daze, E., Gilham, D., Rakai, B.D., Stotz, S.C., Tsujikawa, L.M., Sarsons, C.D., Studer, D., Rinker, K.D., Jahagirdar, R., Wong, N.C.W., Sweeney, M., Johansson, J.O., Kulikowski, E., 2023. The BET inhibitor apabetalone decreases neuroendothelial proinflammatory activation in vitro and in a mouse model of systemic inflammation. Transl Neurosci 14, 20220332. doi: 10.1515/tnsci-2022-0332.

Wasiak, S., Gilham, D., Tsujikawa, L.M., Halliday, C., Calosing, C., Jahagirdar, R., Johansson, J., Sweeney, M., Wong, N.C., Kulikowski, E., 2017. Downregulation of the Complement Cascade In Vitro, in Mice and in Patients with Cardiovascular Disease by the BET Protein Inhibitor Apabetalone (RVX-208). J Cardiovasc. Transl. Res. 10, 337–347. doi: 10.1007/s12265-017-9755-z.

Weinstock, M., 2024. Therapeutic agents for Alzheimer’s disease: a critical appraisal. Front. Aging Neurosci. 16, 1484615. doi: 10.3389/fnagi.2024.1484615.

Wienerroither, S., Rauch, I., Rosebrock, F., Jamieson, A.M., Bradner, J., Muhar, M., Zuber, J., Müller, M., Decker, T., 2014. Regulation of NO synthesis, local inflammation, and innate immunity to pathogens by BET family proteins. Mol. Cell. Biol. 34, 415–427. doi: 10.1128/mcb.01353-13.

Yang, Y.M., Shi, R.H., Xu, C.X., Li, L., 2018. BRD4 expression in patients with essential hypertension and its effect on blood pressure in spontaneously hypertensive rats. J. Am. Soc. Hypertens. 12, e107–e117. doi: 10.1016/j.jash.2018.11.004.

