## Supplementary material for "BET protein inhibitor JQ1 reduces inflammation and hippocampal amyloid-β level without altering Tau phosphorylation in LPS-challenged adult wild-type mice"

Research Article

Matuszewska et al.

### Supplementary Material

#### Supplementary Figures

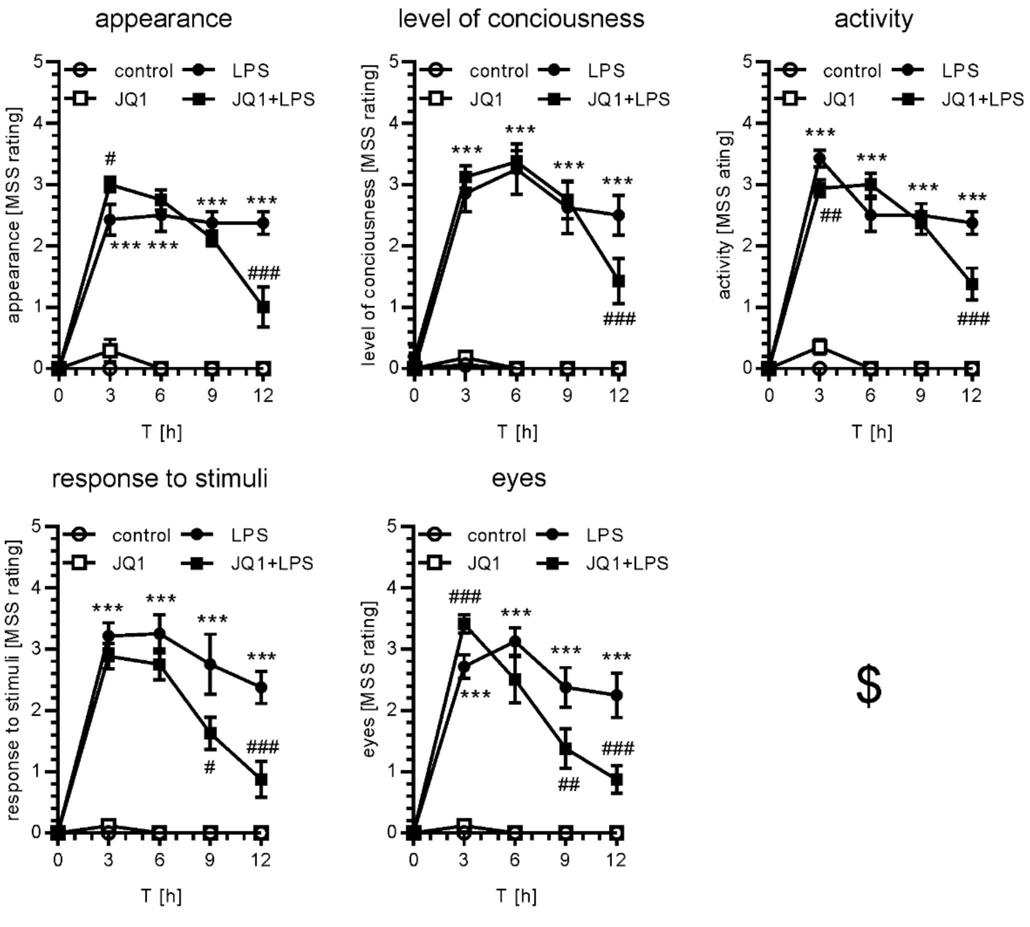


**Supplementary Figure 1:** The effect of LPS and JQ1 on behavioral components of the Murine Sepsis Score (MSS). $ The respiration rate was not analyzed. Mean±SEM was presented. *** $p<0.001$ compared to corresponding time point control; #, ##, ### $p<0.05$, 0.01, and 0.001, respectively, compared to corresponding LPS group. n=8-17. Statistical analyses were conducted using one-way ANOVA followed by the Bonferroni post hoc test.

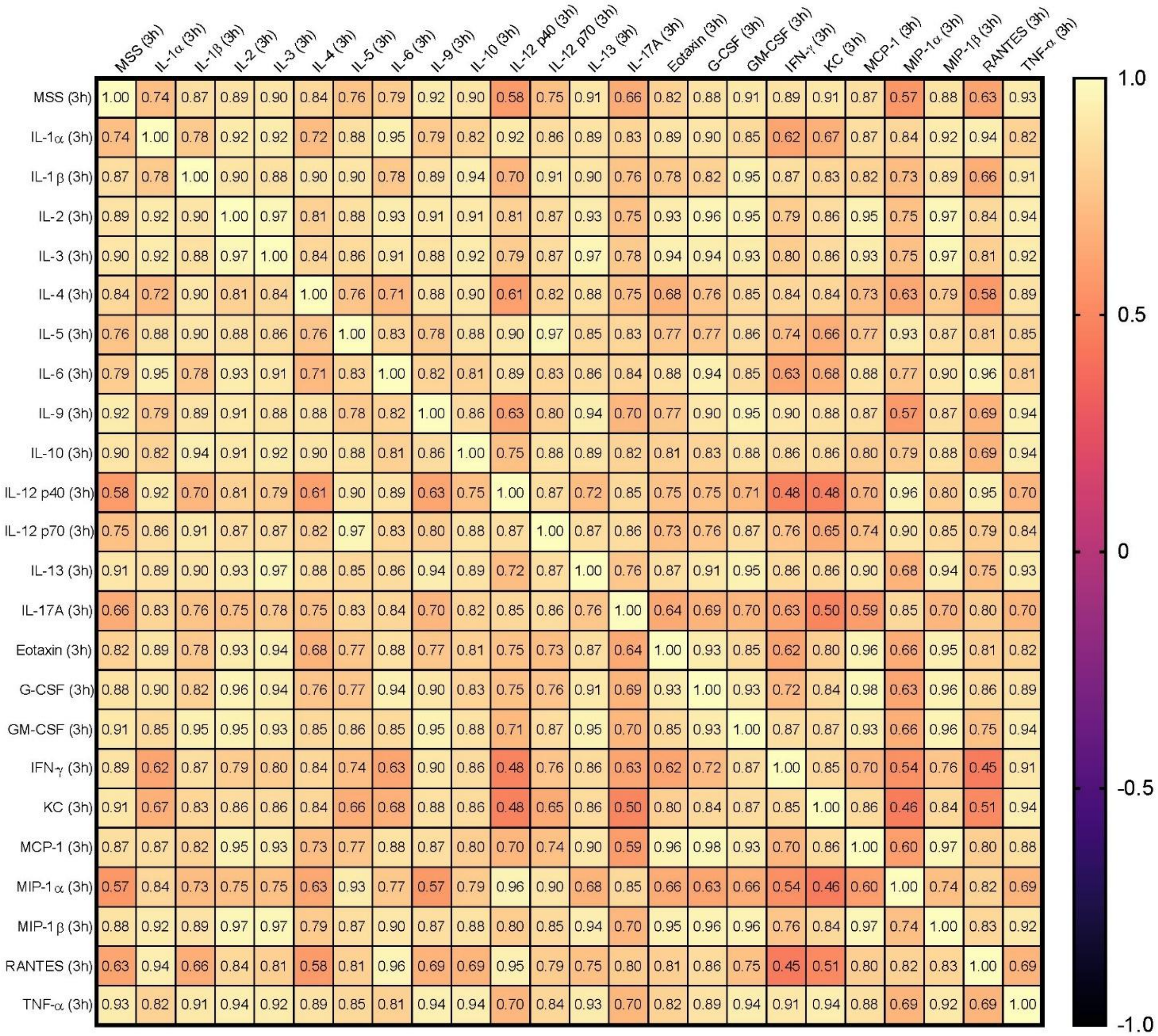


**Supplementary Figure 2a:** The correlation matrix of MSS and blood serum cytokines levels in all experimental groups 3 post-injection. Correlation coefficient R is presented. The analysis of the correlation was performed using Pearson’s correlation test.

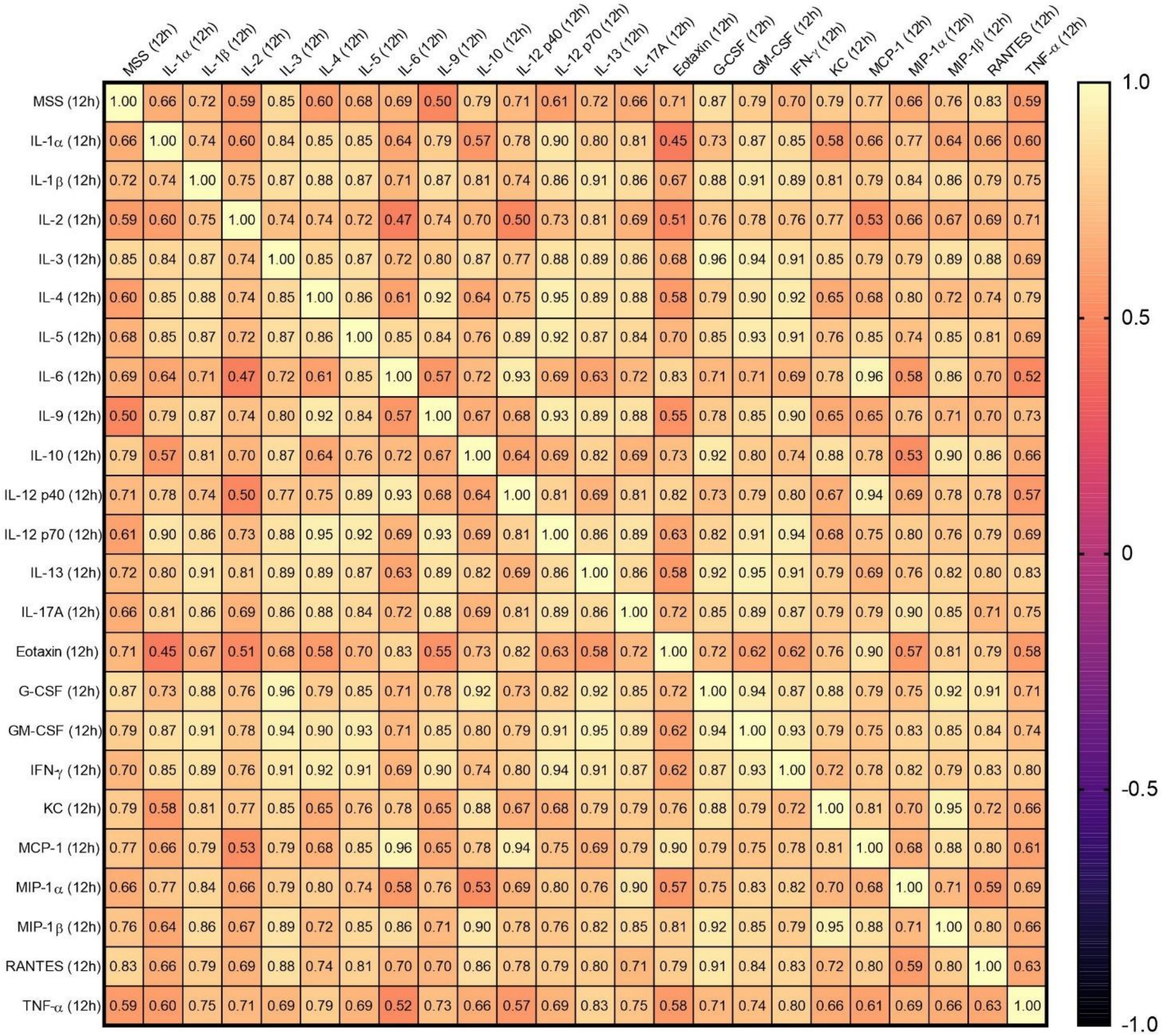


| | MSS (12h) | IL-1α (12h) | IL-1β (12h) | IL-2 (12h) | IL-3 (12h) | IL-4 (12h) | IL-5 (12h) | IL-6 (12h) | IL-9 (12h) | IL-10 (12h) | IL-12 p40 (12h) | IL-12 p70 (12h) | IL-13 (12h) | IL-17A (12h) | Eotaxin (12h) | G-CSF (12h) | GM-CSF (12h) | IFN-γ (12h) | KC (12h) | MCP-1 (12h) | MIP-1α (12h) | MIP-1β (12h) | RANTES (12h) | TNF-α (12h) |
|---|---|---|---|---|---|---|---|---|---|---|---|---|---|---|---|---|---|---|---|---|---|---|---|---|
| MSS (12h) | 1.00 | 0.66 | 0.72 | 0.59 | 0.85 | 0.60 | 0.68 | 0.69 | 0.50 | 0.79 | 0.71 | 0.61 | 0.72 | 0.66 | 0.71 | 0.87 | 0.79 | 0.70 | 0.79 | 0.77 | 0.66 | 0.76 | 0.83 | 0.59 |
| IL-1α (12h) | 0.66 | 1.00 | 0.74 | 0.60 | 0.84 | 0.85 | 0.85 | 0.64 | 0.79 | 0.57 | 0.78 | 0.90 | 0.80 | 0.81 | 0.45 | 0.73 | 0.87 | 0.85 | 0.58 | 0.66 | 0.77 | 0.64 | 0.66 | 0.60 |
| IL-1β (12h) | 0.72 | 0.74 | 1.00 | 0.75 | 0.87 | 0.88 | 0.87 | 0.71 | 0.87 | 0.81 | 0.74 | 0.86 | 0.91 | 0.86 | 0.67 | 0.88 | 0.91 | 0.89 | 0.81 | 0.79 | 0.84 | 0.86 | 0.79 | 0.75 |
| IL-2 (12h) | 0.59 | 0.60 | 0.75 | 1.00 | 0.74 | 0.74 | 0.72 | 0.47 | 0.74 | 0.70 | 0.50 | 0.73 | 0.81 | 0.69 | 0.51 | 0.76 | 0.78 | 0.76 | 0.77 | 0.53 | 0.66 | 0.67 | 0.69 | 0.71 |
| IL-3 (12h) | 0.85 | 0.84 | 0.87 | 0.74 | 1.00 | 0.85 | 0.87 | 0.72 | 0.80 | 0.87 | 0.77 | 0.88 | 0.89 | 0.86 | 0.68 | 0.96 | 0.94 | 0.91 | 0.85 | 0.79 | 0.79 | 0.89 | 0.88 | 0.69 |
| IL-4 (12h) | 0.60 | 0.85 | 0.88 | 0.74 | 0.85 | 1.00 | 0.86 | 0.61 | 0.92 | 0.64 | 0.75 | 0.95 | 0.89 | 0.88 | 0.58 | 0.79 | 0.90 | 0.92 | 0.65 | 0.68 | 0.80 | 0.72 | 0.74 | 0.79 |
| IL-5 (12h) | 0.68 | 0.85 | 0.87 | 0.72 | 0.87 | 0.86 | 1.00 | 0.85 | 0.84 | 0.76 | 0.89 | 0.92 | 0.87 | 0.84 | 0.70 | 0.85 | 0.93 | 0.91 | 0.76 | 0.85 | 0.74 | 0.85 | 0.81 | 0.69 |
| IL-6 (12h) | 0.69 | 0.64 | 0.71 | 0.47 | 0.72 | 0.61 | 0.85 | 1.00 | 0.57 | 0.72 | 0.93 | 0.69 | 0.63 | 0.72 | 0.83 | 0.71 | 0.71 | 0.69 | 0.78 | 0.96 | 0.58 | 0.86 | 0.70 | 0.52 |
| IL-9 (12h) | 0.50 | 0.79 | 0.87 | 0.74 | 0.80 | 0.92 | 0.84 | 0.57 | 1.00 | 0.67 | 0.68 | 0.93 | 0.89 | 0.88 | 0.55 | 0.78 | 0.85 | 0.90 | 0.65 | 0.65 | 0.76 | 0.71 | 0.70 | 0.73 |
| IL-10 (12h) | 0.79 | 0.57 | 0.81 | 0.70 | 0.87 | 0.64 | 0.76 | 0.72 | 0.67 | 1.00 | 0.64 | 0.69 | 0.82 | 0.69 | 0.73 | 0.92 | 0.80 | 0.74 | 0.88 | 0.78 | 0.53 | 0.90 | 0.86 | 0.66 |
| IL-12 p40 (12h) | 0.71 | 0.78 | 0.74 | 0.50 | 0.77 | 0.75 | 0.89 | 0.93 | 0.68 | 0.64 | 1.00 | 0.81 | 0.69 | 0.81 | 0.82 | 0.73 | 0.79 | 0.80 | 0.67 | 0.94 | 0.69 | 0.78 | 0.78 | 0.57 |
| IL-12 p70 (12h) | 0.61 | 0.90 | 0.86 | 0.73 | 0.88 | 0.95 | 0.92 | 0.69 | 0.93 | 0.69 | 0.81 | 1.00 | 0.86 | 0.89 | 0.63 | 0.82 | 0.91 | 0.94 | 0.68 | 0.75 | 0.80 | 0.76 | 0.79 | 0.69 |
| IL-13 (12h) | 0.72 | 0.80 | 0.91 | 0.81 | 0.89 | 0.89 | 0.87 | 0.63 | 0.89 | 0.82 | 0.69 | 0.86 | 1.00 | 0.86 | 0.58 | 0.92 | 0.95 | 0.91 | 0.79 | 0.69 | 0.76 | 0.82 | 0.80 | 0.83 |
| IL-17A (12h) | 0.66 | 0.81 | 0.86 | 0.69 | 0.86 | 0.88 | 0.84 | 0.72 | 0.88 | 0.69 | 0.81 | 0.89 | 0.86 | 1.00 | 0.72 | 0.85 | 0.89 | 0.87 | 0.79 | 0.79 | 0.90 | 0.85 | 0.71 | 0.75 |
| Eotaxin (12h) | 0.71 | 0.45 | 0.67 | 0.51 | 0.68 | 0.58 | 0.70 | 0.83 | 0.55 | 0.73 | 0.82 | 0.63 | 0.58 | 0.72 | 1.00 | 0.72 | 0.62 | 0.62 | 0.76 | 0.90 | 0.57 | 0.81 | 0.79 | 0.58 |
| G-CSF (12h) | 0.87 | 0.73 | 0.88 | 0.76 | 0.96 | 0.79 | 0.85 | 0.71 | 0.78 | 0.92 | 0.73 | 0.82 | 0.92 | 0.85 | 0.72 | 1.00 | 0.94 | 0.87 | 0.88 | 0.79 | 0.75 | 0.92 | 0.91 | 0.71 |
| GM-CSF (12h) | 0.79 | 0.87 | 0.91 | 0.78 | 0.94 | 0.90 | 0.93 | 0.71 | 0.85 | 0.80 | 0.79 | 0.91 | 0.95 | 0.89 | 0.62 | 0.94 | 1.00 | 0.93 | 0.79 | 0.75 | 0.83 | 0.85 | 0.84 | 0.74 |
| IFN-γ (12h) | 0.70 | 0.85 | 0.89 | 0.76 | 0.91 | 0.92 | 0.91 | 0.69 | 0.90 | 0.74 | 0.80 | 0.94 | 0.91 | 0.87 | 0.62 | 0.87 | 0.93 | 1.00 | 0.72 | 0.78 | 0.82 | 0.79 | 0.83 | 0.80 |
| KC (12h) | 0.79 | 0.58 | 0.81 | 0.77 | 0.85 | 0.65 | 0.76 | 0.78 | 0.65 | 0.88 | 0.67 | 0.68 | 0.79 | 0.79 | 0.76 | 0.88 | 0.79 | 0.72 | 1.00 | 0.81 | 0.70 | 0.95 | 0.72 | 0.66 |
| MCP-1 (12h) | 0.77 | 0.66 | 0.79 | 0.53 | 0.79 | 0.68 | 0.85 | 0.96 | 0.65 | 0.78 | 0.94 | 0.75 | 0.69 | 0.79 | 0.90 | 0.79 | 0.75 | 0.78 | 0.81 | 1.00 | 0.68 | 0.88 | 0.80 | 0.61 |
| MIP-1α (12h) | 0.66 | 0.77 | 0.84 | 0.66 | 0.79 | 0.80 | 0.74 | 0.58 | 0.76 | 0.53 | 0.69 | 0.80 | 0.76 | 0.90 | 0.57 | 0.75 | 0.83 | 0.82 | 0.70 | 0.68 | 1.00 | 0.71 | 0.59 | 0.69 |
| MIP-1β (12h) | 0.76 | 0.64 | 0.86 | 0.67 | 0.89 | 0.72 | 0.85 | 0.86 | 0.71 | 0.90 | 0.78 | 0.76 | 0.82 | 0.85 | 0.81 | 0.92 | 0.85 | 0.79 | 0.95 | 0.88 | 0.71 | 1.00 | 0.80 | 0.66 |
| RANTES (12h) | 0.83 | 0.66 | 0.79 | 0.69 | 0.88 | 0.74 | 0.81 | 0.70 | 0.70 | 0.86 | 0.78 | 0.79 | 0.80 | 0.71 | 0.79 | 0.91 | 0.84 | 0.83 | 0.72 | 0.80 | 0.59 | 0.80 | 1.00 | 0.63 |
| TNF-α (12h) | 0.59 | 0.60 | 0.75 | 0.71 | 0.69 | 0.79 | 0.69 | 0.52 | 0.73 | 0.66 | 0.57 | 0.69 | 0.83 | 0.75 | 0.58 | 0.71 | 0.74 | 0.80 | 0.66 | 0.61 | 0.69 | 0.66 | 0.63 | 1.00 |

**Supplementary Figure 2b:** The correlation matrix of MSS and blood serum cytokines levels in all experimental groups 12 h post-injection. Correlation coefficient R is presented. The analysis of the correlation was performed using Pearson's correlation test.

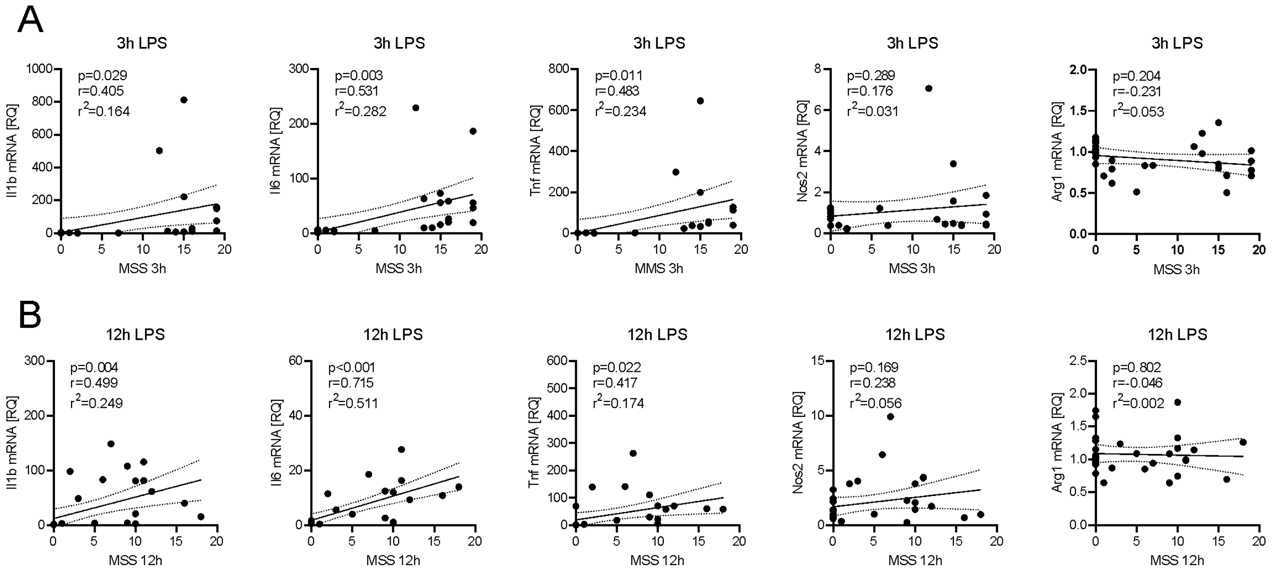


**Supplementary Figure 3:** Correlation of the sickness behavior score (MSS) with inflammatory genes' expression in hippocampus 3 h (A) and 12 h (B) after administration of LPS.

Data presented in the manuscript on Figure 1 and Figure 4 from all experimental groups were utilized to correlation analysis using Pearson's correlation test. The full lines indicate the linear regression, the dashed lines indicate 95% confidence bands.

Supplementary Tables

**Supplementary Table 1**

| **Experimental conditions used to perform the Western blot experiments presented on Figure 5.** | | |
|---|---|---|
| **Primary antibody** | **Brand / Cat #** | **Dilution** |
| mouse anti-Tau IgG | Santa Cruz Biotechnology<br>sc-32274 | 1:500<br>5% milk in TBS-T 0.1% |
| rabbit anti-phospho-Tau IgG<br>(Ser199-202) | Sigma-Aldrich<br>T6819 | 1:1000<br>5% milk in TBS-T 0.1% |
| mouse anti-phospho-Tau IgG<br>(Ser396) | Cell Signalling Technology<br>9632 | 1:250<br>TBS-T 0.1% |
| rabbit anti-phospho-Tau IgG<br>(Ser416) | Cell Signalling Technology<br>15013 | 1:1000<br>5% milk in TBS-T 0.1% |
| rabbit anti-GAPDH IgG | Sigma-Aldrich<br>G9545 | 1:40,000<br>5% milk in TBS-T 0.1% |
| **Secondary antibody** | | |
| goat anti-mouse IgG (HRP-conjugated) | Thermo Fisher Scientific<br>A28177 | 1:4000<br>5% milk in TBS-T 0.1% |
| goat anti-rabbit IgG (HRP-conjugated) | Sigma-Aldrich<br>A0545 | 1:4000<br>5% milk in TBS-T 0.1% |

#### Supplementary Table 2

| | control | | | 3h LPS | | | 3h LPS+JQ1 | | | 3h JQ1 | | | 12h LPS | | | 12h LPS+JQ1 | | | 12h JQ1 | | |
|---|---|---|---|---|---|---|---|---|---|---|---|---|---|---|---|---|---|---|---|---|---|
| | mean± | SEM | N | mean± | SEM | N | mean± | SEM | N | mean± | SEM | N | mean± | SEM | N | mean± | SEM | N | mean± | SEM | N |
| IL-1α [pg/mL] | 18.99± | 2.075 | 7 | 64.94± | 12.38 | 5 *** | 33.68± | 1.343 | 4 # | 15.14± | 0.454 | 3 | 51.73± | 13.91 | 5 * | 32.68± | 5.58 | 5 | 18.16± | 10.09 | 4 |
| IL-1β [pg/mL] | 13.52± | 2.603 | 8 | 46.11± | 9.91 | 5 ** | 34.5± | 2.057 | 4 | 15.02± | 3.304 | 4 | 41.39± | 5.28 | 5 * | 37.15± | 10.29 | 6 | 10.62± | 0.87 | 4 |
| IL-2 [pg/mL] | 22.25± | 1.384 | 8 | 86.12± | 13.22 | 5 *** | 57.34± | 2.485 | 4 P<0.1 | 31.41± | 8.845 | 4 | 85.42± | 2.71 | 4 * | 85.53± | 28.68 | 6 | 14.24± | 4.95 | 4 |
| IL-3 [pg/mL] | 2.451± | 0.841 | 8 | 36.56± | 10.49 | 5 *** | 31.06± | 0.767 | 4 | 1.84± | 0.402 | 4 | 51.06± | 1.46 | 4 *** | 35.31± | 10.06 | 6 | 0.89± | 0.12 | 3 |
| IL-4 [pg/mL] | 8.31± | 1.47 | 8 | 25.62± | 6.45 | 5 ** | 19.73± | 1.846 | 4 | 10.71± | 3.524 | 4 | 21.82± | 4.08 | 5 * | 19.95± | 5.26 | 6 | 3.98± | 0.14 | 3 |
| IL-5 [pg/mL] | 22.28± | 2.268 | 8 | 107.90± | 29.54 | 5 ** | 54.97± | 2.49 | 4 | 24.45± | 0.799 | 3 | 79.08± | 15.17 | 5 *** | 44.59± | 9.77 | 5 # | 22.71± | 1.78 | 4 |
| IL-6 [pg/mL] | 25.61± | 10.8 | 9 | 12825.0± | 4784.00 | 5 *** | 6415± | 252.2 | 4 | 58.49± | 8.011 | 4 | 2646.00± | 994.30 | 5 *** | 478.30± | 144.10 | 6 ## | 9.81± | 3.35 | 4 |
| IL-9 [pg/mL] | 112.9± | 7.84 | 8 | 197.30± | 20.88 | 4 *** | 192.1± | 6.486 | 4 | 110.5± | 6.943 | 3 | 192.00± | 23.87 | 5 | 208.30± | 73.20 | 6 | 103.40± | 10.42 | 4 |
| IL-10 [pg/mL] | 170± | 27.13 | 8 | 2081.00± | 634.50 | 5 ** | 2365± | 334.7 | 4 | 405.8± | 126 | 4 | 759.20± | 181.10 | 5 * | 644.20± | 174.80 | 6 | 132.20± | 46.49 | 4 |
| IL-12p40 [pg/mL] | 460.6± | 132.5 | 9 | 7445.00± | 3872.00 | 5 * | 2274± | 305.1 | 4 | 245.9± | 31.8 | 4 | 5962.00± | 1950.00 | 5 *** | 727.10± | 220.30 | 6 ### | 190.10± | 85.55 | 4 |
| IL-12p70 [pg/mL] | 71.81± | 4.417 | 8 | 323.00± | 82.70 | 5 *** | 178.4± | 4.609 | 4 P<0.1 | 100.3± | 29.83 | 4 | 219.80± | 36.15 | 5 * | 186.90± | 65.09 | 6 | 91.40± | 24.97 | 4 |
| IL-13 [pg/mL] | 71.13± | 15.56 | 8 | 354.20± | 84.65 | 5 *** | 242.1± | 1.59 | 4 | 66.48± | 15.75 | 4 | 213.30± | 35.31 | 5 ** | 175.20± | 35.79 | 5 | 33.79± | 5.86 | 4 |
| IL-17A [pg/mL] | 62.97± | 3.74 | 8 | 162.60± | 52.35 | 5 * | 85.29± | 6.623 | 4 | 73.43± | 14.58 | 4 | 156.80± | 29.86 | 5 * | 134.50± | 34.89 | 6 | 48.19± | 1.26 | 3 |
| Eotaxin [pg/mL] | 4297± | 201.1 | 9 | 8229.00± | 2839.00 | 5 P<0.1 | 9164± | 559.8 | 4 | 3700± | 168.4 | 4 | 9695.00± | 1006.00 | 5 *** | 4805.00± | 663.20 | 6 ### | 4585.00± | 467.50 | 4 |
| G-CSF [pg/mL] | 1619± | 1010 | 9 | 80409.0± | 17666.0 | 5 *** | 55858± | 459.8 | 3 | 1233± | 175.2 | 4 | 157644.0± | 7403.00 | 4 *** | 104646.00± | 28984.0 | 6 | 365.90± | 102.80 | 4 |
| GM-CSF [pg/mL] | 143.8± | 8.465 | 8 | 407.10± | 6.74 | 3 *** | 332.5± | 4.878 | 4 ### | 147.8± | 3.15 | 3 | 362.10± | 52.37 | 5 *** | 290.00± | 42.91 | 5 | 138.00± | 6.16 | 4 |
| IFN-γ [pg/mL] | 67.73± | 7.131 | 8 | 167.90± | 35.95 | 4 ** | 198.1± | 1.021 | 3 | 101.5± | 23.78 | 4 | 192.80± | 45.86 | 5 P<0.1 | 185.40± | 59.37 | 6 | 62.59± | 10.81 | 4 |
| KC [pg/mL] | 90.43± | 7.918 | 7 | 8395.00± | 1878.00 | 5 *** | 10628± | 60.97 | 3 | 356± | 95.92 | 4 | 7825.00± | 2721.00 | 5 ** | 5847.00± | 1709.00 | 6 | 103.00± | 9.07 | 4 |
| MCP-1 [pg/mL] | 264.7± | 44.83 | 7 | 36445.0± | 14227.0 | 5 ** | 20594± | 430.1 | 4 | 506.9± | 75.71 | 4 | 11881.00± | 3370.00 | 5 *** | 1861.00± | 469.80 | 6 ### | 330.50± | 73.87 | 4 |
| MIP-1α [pg/mL] | 7.208± | 1.621 | 9 | 1021.00± | 467.70 | 5 ** | 278± | 31.66 | 4 | 10.61± | 0.65 | 3 | 74.51± | 29.38 | 5 * | 59.97± | 15.72 | 6 | 10.31± | 0.34 | 3 |
| MIP-1β [pg/mL] | 121.1± | 32.91 | 7 | 9488.00± | 1953.00 | 5 *** | 7541± | 364.3 | 4 | 204.1± | 43.45 | 4 | 1373.00± | 387.00 | 5 ** | 860.50± | 212.50 | 6 | 105.40± | 10.95 | 4 |
| RANTES [pg/mL] | 400.7± | 65.21 | 9 | 15164.0± | 4647.00 | 5 *** | 3209± | 267 | 4 ## | 112.4± | 5.399 | 4 | 21769.00± | 5729.00 | 5 *** | 8469.00± | 2147.00 | 6 ## | 156.20± | 33.46 | 4 |
| TNF-α [pg/mL] | 104.1± | 10.03 | 8 | 426.00± | 101.00 | 5 ** | 308.2± | 32.6 | 4 | 89.01± | 1.935 | 3 | 320.60± | 57.58 | 5 * | 370.00± | 85.84 | 6 | 80.32± | 12.30 | 4 |

The effect of LPS and JQ1 on the level of pro-inflammatory mediators in blood serum 3 and 12 h post-injection. *,**,*** p<0.05, p<0.01, and p<0.001, compared to control. #, ##, ### p<0.05, p<0.01, and p<0.001, compared to respective LPS-group. Statistical analyses were conducted using one-way ANOVA followed by the Bonferroni post hoc test.

**Supplementary Table 3**

| | **control [FC]** | **JQ1 [FC]** |
|---|---|---|
| *Abca7* | 1.00±0.10 | 2.37±0.17** |
| *Bin1* | 1.00±0.03 | 0.85±0.01* |
| *Cd2ap* | 1.00±0.11 | 0.80±0.04 |
| *Clu* | 1.00±0.11 | 2.59±0.16*** |
| *Cr1l* | 1.00±0.14 | 0.63±0.04 |
| *Picalm* | 1.00±0.07 | 0.86±0.02 |
| *Rin3* | 1.00±0.09 | 0.57±0.05* |
| *Trem2* | 1.00±0.21 | 0.97±0.21 |
| *Zyx* | 1.00±0.11 | 1.09±0.07 |

The curated analysis of the effect of 4 h incubation in the presence of 500 nM JQ1 on expression of phagocytosis/endocytosis-related genes in human microglial cell line HMC3 (Baek et al., 2021).

The curated analysis of the Baek and co-workers used RNA sequencing to analyze the impact of JQ1 on gene expression in human microglial HMC3 line (Baek et al., 2021). The expression data that have been deposited in NCBI's Gene Expression Omnibus (Edgar et al., 2002) and are accessible through GEO Series accession number GSE155408 (https://www.ncbi.nlm.nih.gov/geo/query/acc.cgi?acc=GSE155408). *,**,*** p<0.05, p<0.01, and p<0.001, compared to control. Statistical analyses were conducted using the Student’s t-test. FC – fold-change.

Supplementary file - the full uncropped blots images

**Gapdh**

C - control
L - LPS
LJ - LPS+JQ1
J - JQ1
red frame - samples presented in the manuscript

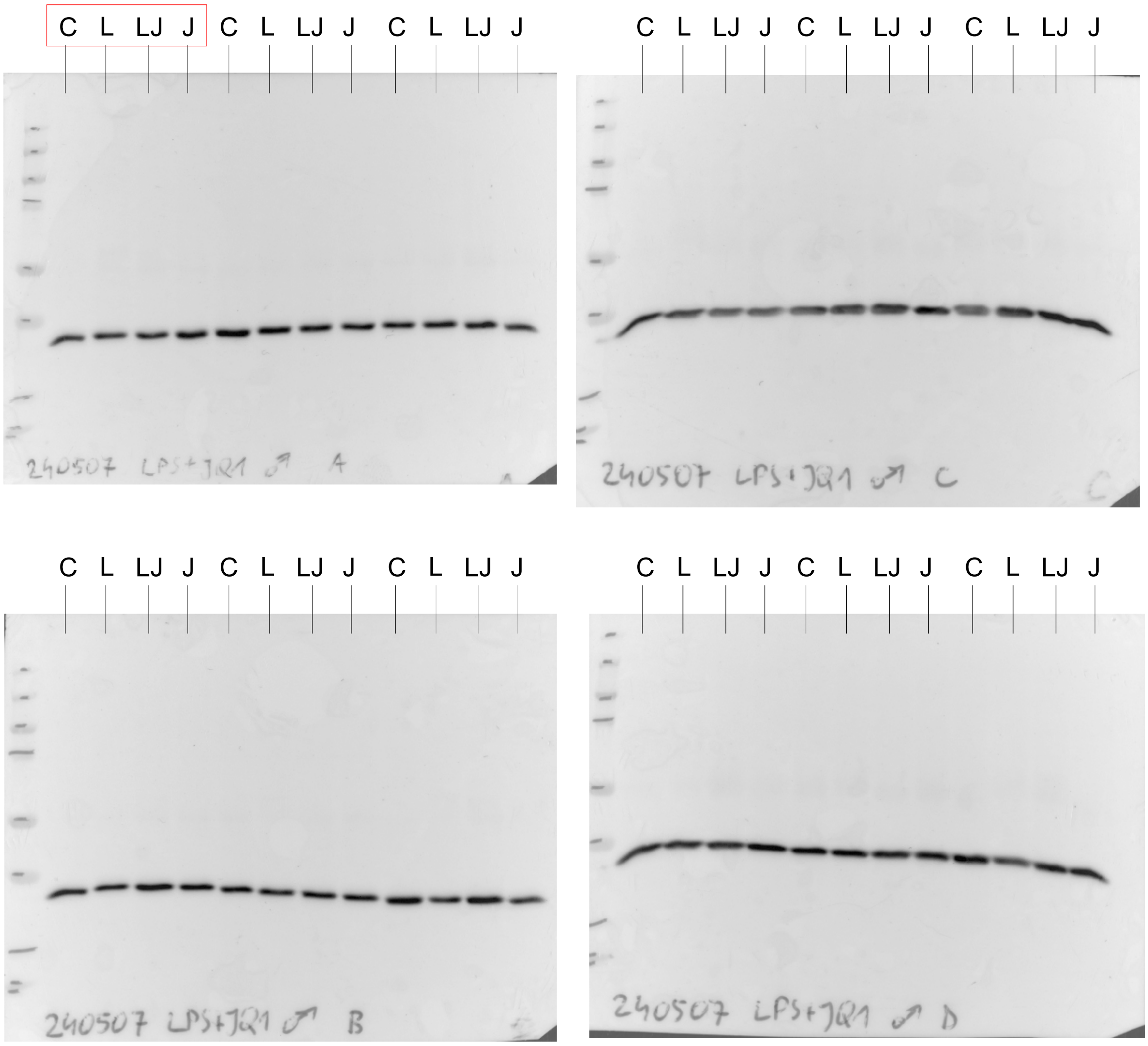


Membranes 240507A and 240507C contain the set of 12 samples.
Membranes 240507B and 240507D contain the other set of 12 samples.

**total Tau**

C - control
L - LPS
LJ - LPS+JQ1
J - JQ1
red frame - samples presented in the manuscript

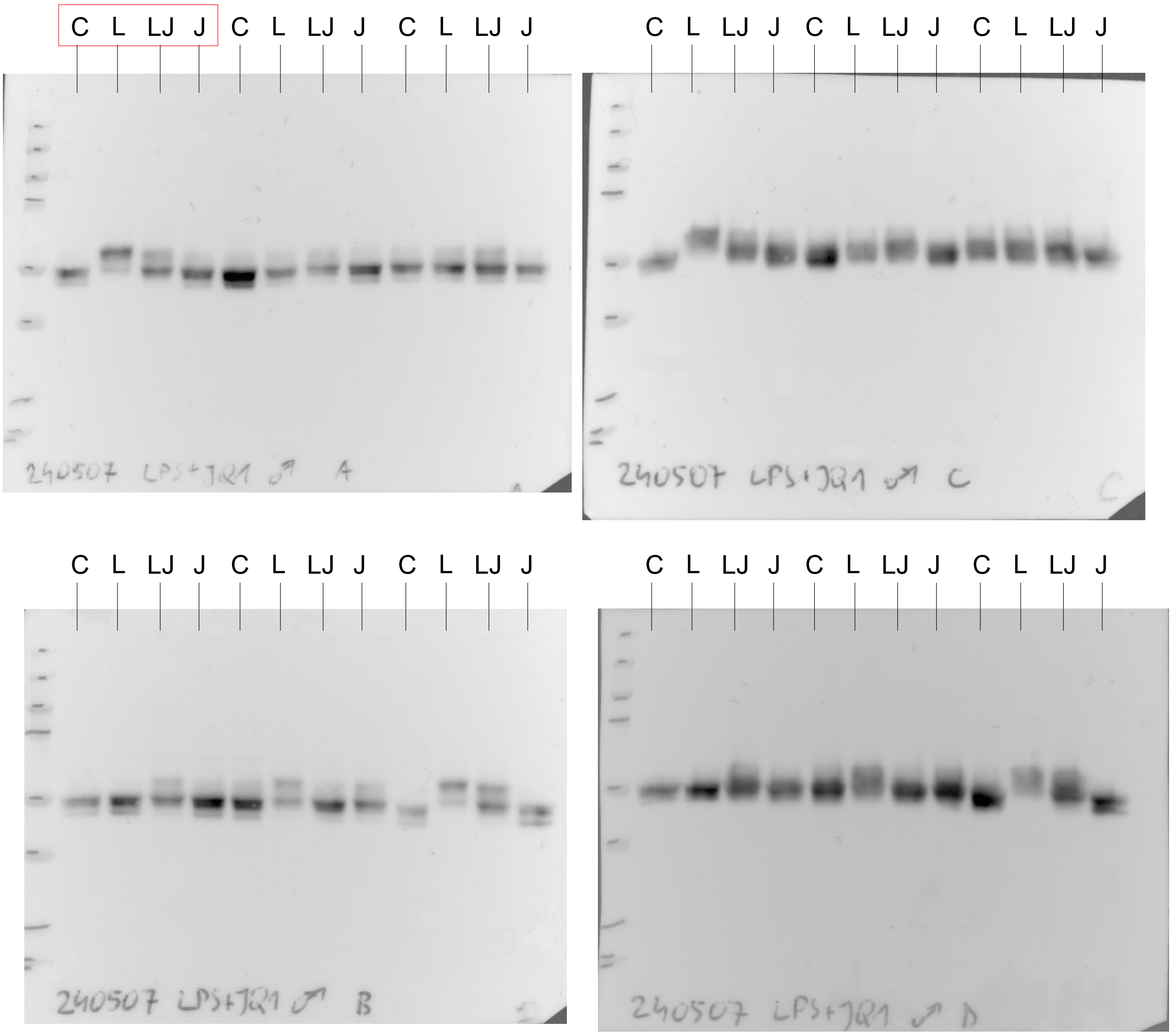


Membranes 240507A and 240507C contain the set of 12 samples.
Membranes 240507B and 240507D contain the other set of 12 samples.

**phospho-Tau (Ser199-202)**

C - control
L - LPS
LJ - LPS+JQ1
J - JQ1
red frame - samples presented in the manuscript

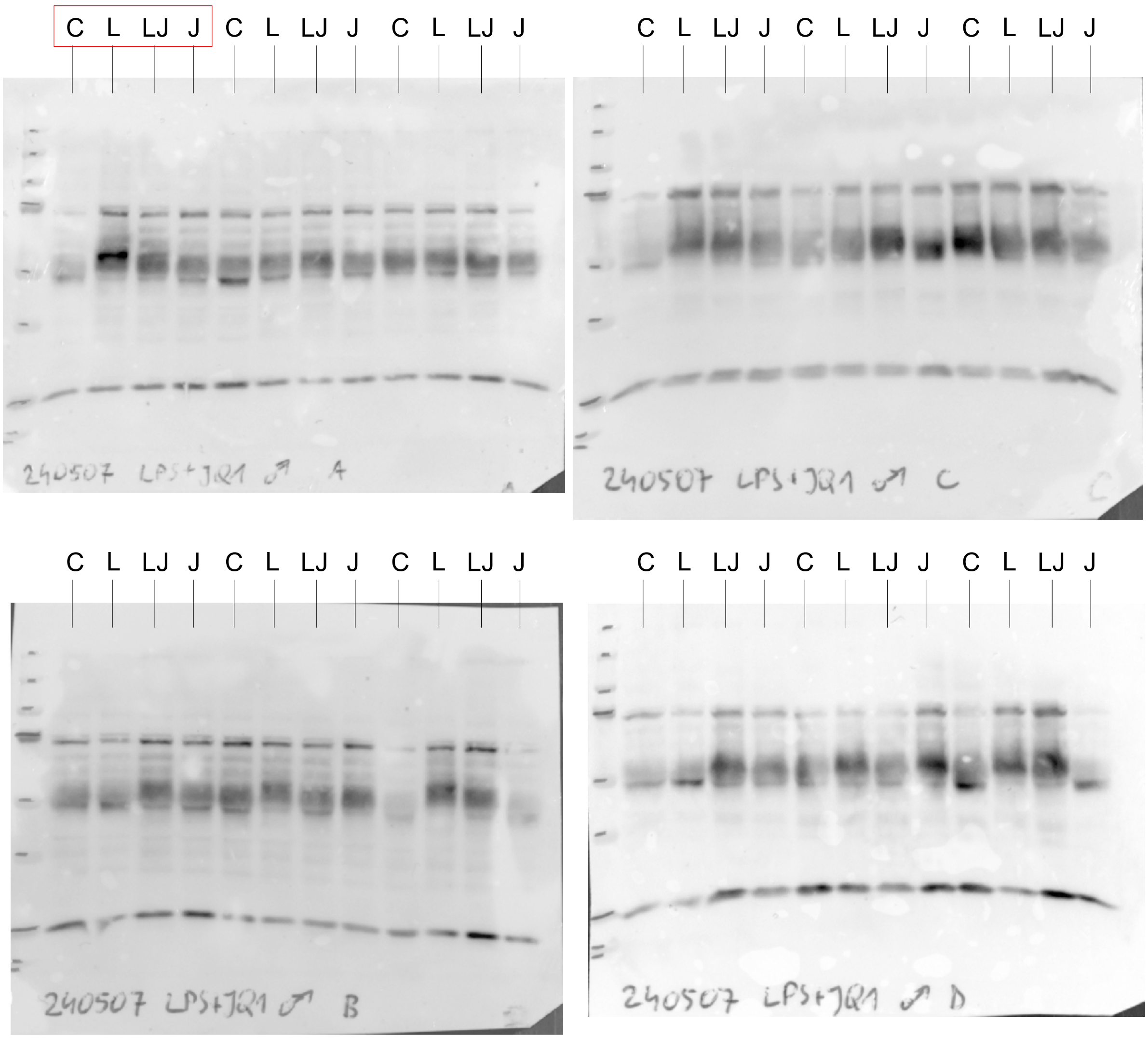


Membranes 240507A and 240507C contain the set of 12 samples.
Membranes 240507B and 240507D contain the other set of 12 samples.

**phospho-Tau (Ser396)**

C - control
L - LPS
LJ - LPS+JQ1
J - JQ1
red frame - samples presented in the manuscript

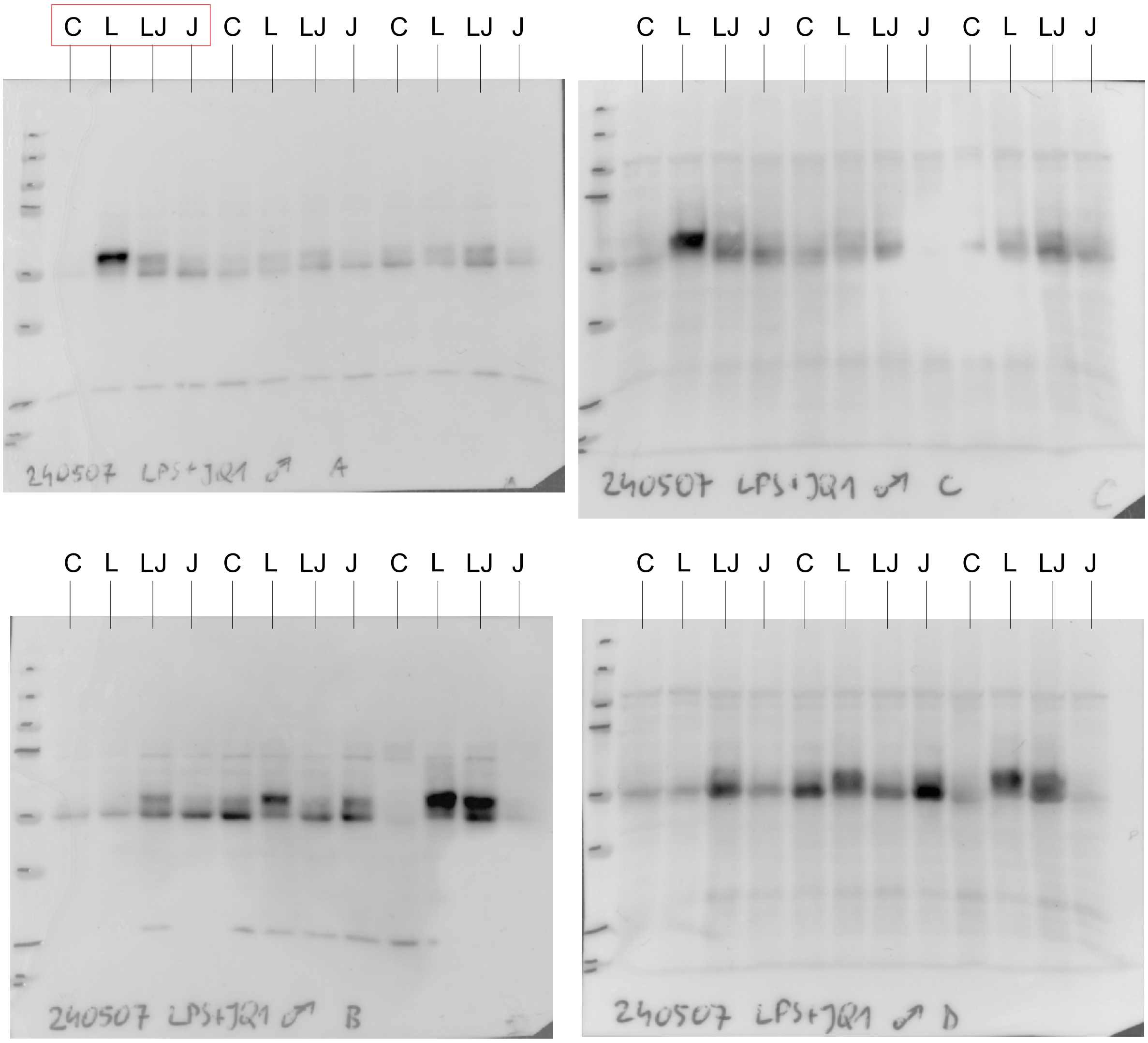


Membranes 240507A and 240507C contain the set of 12 samples.
Membranes 240507B and 240507D contain the other set of 12 samples.

**phospho-Tau (Ser416)**

C - control
L - LPS
LJ - LPS+JQ1
J - JQ1
red frame - samples presented in the manuscript

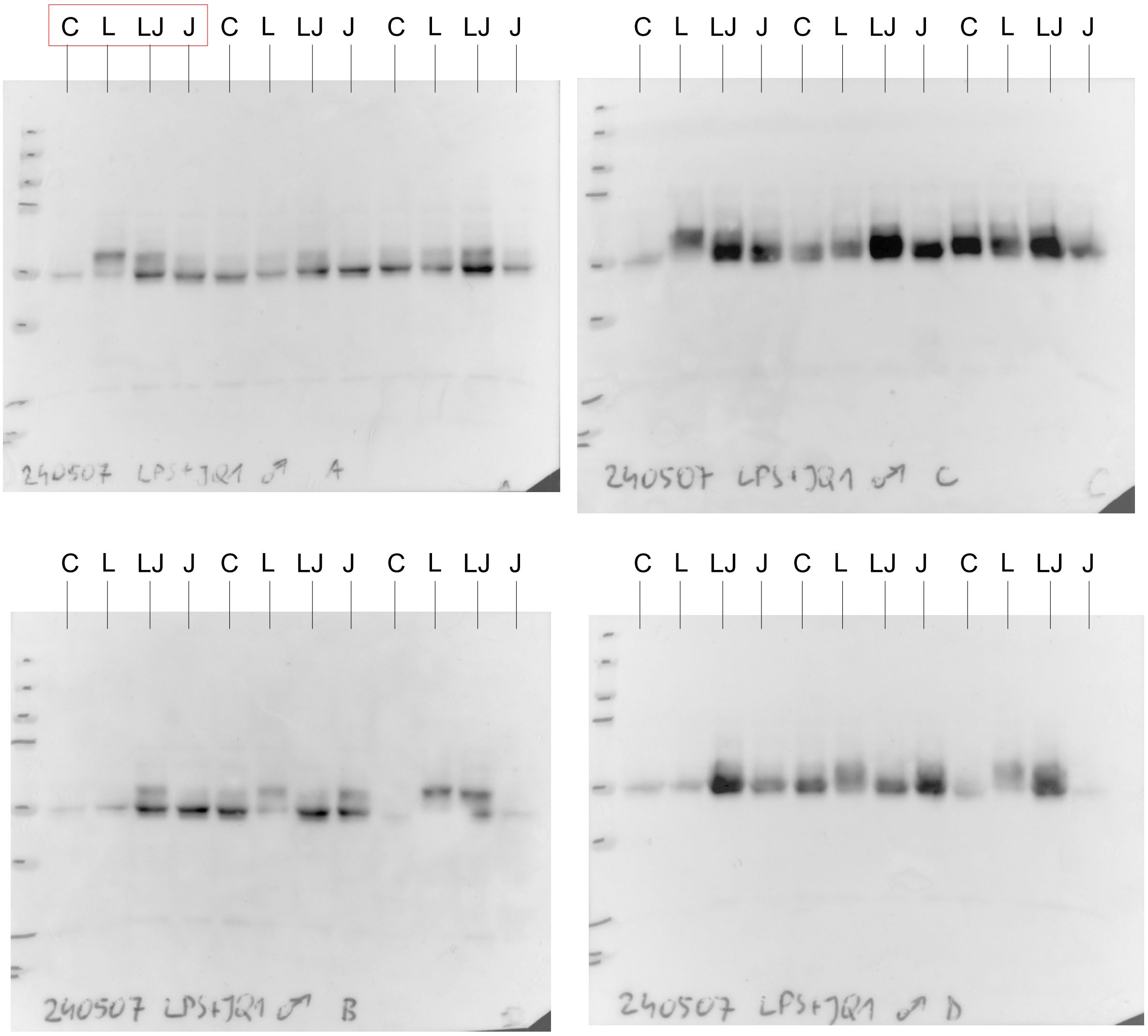


Membranes 240507A and 240507C contain the set of 12 samples.
Membranes 240507B and 240507D contain the other set of 12 samples.
